# Loss of the alternative calcineurin variant CnAβ1 enhances brown adipocyte differentiation and drives metabolic overactivation through FoxO1 activation

**DOI:** 10.1101/2025.05.09.653054

**Authors:** Elísabet Bello-Arroyo, Blanca Rubio, Alfonso Mora, Marina López-Olañeta, Jesús M. Gómez-Salinero, Laura Ramos, Carlos G. Sánchez-Cabezudo, Emilio Camafeita, Lorena Cusso, Manuel Desco, Nolwenn Joffin, Johanne Le Coq, Jasminka Boskovic, Kathryn A. McGurk, James S. Ware, Paul J. R. Barton, Jesús Vázquez, Philipp E. Scherer, Guadalupe Sabio, María Victoria Gómez-Gaviro, Enrique Lara-Pezzi

## Abstract

The alternative calcineurin A variant CnAβ1 has a unique C-terminal domain that provides it with distinct subcellular localization and mechanism of action different from other calcineurin isoforms. Here, we used mice lacking CnAβ1’s C-terminal domain (CnAβ1^Δi12^) to show that the absence of this specific isoform strongly reprograms metabolism. CnAβ1^Δi12^ mice on a high-fat diet showed reduced body weight, white adipose tissue (WAT) mass, and circulating triglycerides, together with enhanced insulin sensitivity. In brown adipose tissue (BAT), CnAβ1 deficiency increased mitochondrial content and upregulated fatty acid oxidation and thermogenic proteins, improving cold resistance. Conversely, under starvation, CnAβ1^Δi12^ mice experienced rapid fat depletion and hypothermia. Importantly, BAT-specific FoxO1 knockout in CnAβ1^Δi12^ mice reduced catabolism-related gene expression and partially reversed the metabolic phenotypes, increasing body weight and WAT mass. Our findings reveal a relevant role for CnAβ1 in orchestrating BAT metabolism, highlighting its potential as a therapeutic target for obesity and metabolic syndrome.

## INTRODUCTION

White adipose tissue (WAT) is the main reservoir of triglycerides (TGs), which are stored in white adipocytes during the postprandial phase^1^. In response to nutrient shortage, fasting, or prolonged cold exposure, TGs are sequentially broken down into glycerol and free fatty acids (FFAs) that are released into the bloodstream and serve as fuel for various organs, including the brown adipose tissue (BAT)^1,2,3^. Unlike white adipocytes, brown adipocytes are multilocular, containing multiple lipid droplets and a high number of mitochondria. In response to cold or high-fat diet, the adrenal gland and the sympathetic nerves release norepinephrine, which stimulates the β3-adrenergic receptor in BAT to increase body temperature through non-shivering thermogenesis (NST)^3^. This process primarily uses FFA as fuel, although the BAT can also use alternative sources of energy like branched-chain amino acids (BCAA)^4^.

The extensive use of FFAs by the BAT leads to the depletion of lipid droplets in both WAT and BAT and contributes to a decrease in body weight^5^. Similarly, BCAA catabolism in the BAT has beneficial metabolic effects on the organism beyond heat generation^6^. Given that body mass index inversely correlates with BAT activity, enhancing the energy-dissipating function of BAT has been proposed as a potential therapeutic strategy for the treatment of obesity. Moreover, recent studies have linked increased BAT activity with improved cardiometabolic health^7^. Therefore, the BAT is considered a major metabolic regulator beyond thermogenesis, contributing to glucose and lipid homeostasis, and communicating with other organs to maintain the metabolic balance in the organism.

Calcineurin (Cn) is a calcium/calmodulin-dependent serine/threonine phosphatase involved in various biological processes, including cardiac hypertrophy and T cell differentiation^8^. Its catalytic subunit, CnA is expressed from three different genes to produce CnAα and CnAβ, which are are ubiquitously expressed, and CnAγ, which is restricted to testis and brain^8,9^. CnAβ is further regulated through alternative polyadenylation to produce two variants with opposing functions: CnAβ2, which is the predominant form and functions like other CnA isoforms, and CnAβ1, which has a unique C-terminal domain that confers distinct properties to this variant^10^.

CnAβ1’s function and mechanism of action are different from those of other CnA isoforms. In skeletal muscle, we previously demonstrated that CnAβ1 enhances muscle regeneration^11^. In the heart, we showed that CnAβ1 overexpression has beneficial effects following myocardial infarction in mice^12,13^, reducing fibrosis and accelerating the resolution of inflammation by activating the transcription factor ATF4 in an mTOR-dependent manner^13^. Remarkably, unlike other calcineurin A isoforms, which strongly promote hypertrophy, CnAβ1 overexpression in the heart does not activate NFAT and does not induce hypertrophy^12,13^. Instead, CnAβ1 reduces maladaptive hypertrophy and cardiac remodelling after pressure overload by activating serine and one-carbon metabolism in a mTOR-dependent manner^14^.

We previously showed that CnAβ is post-transcriptionally regulated during myoblast differentiation, with a progressive reduction of the CnAβ1/CnAβ2 ratio^15^. We also unveiled that loss of CnAβ1 resulted in accelerated differentiation. Since skeletal muscle and BAT share common developmental precursors^16^, we hypothesised that CnAβ1 could regulate brown adipocyte differentiation and BAT function.

In this work, we found that the CnAβ1/CnAβ2 ratio decreased during brown pre-adipocyte differentiation and that loss of CnAβ1 accelerated differentiation. Mice lacking CnAβ1 showed significantly increased mitochondrial content, lipid catabolism and thermogenesis in BAT. This enhanced BAT metabolism led to a reduction in body weight due to decreased fat accumulation in white adipocytes. In the absence of CnAβ1, BAT showed reduced phosphorylation of the mTOR-Akt axis and increased FoxO1 activation. A BAT-specific FoxO1 knockout prevented the activation of lipid catabolism and partially reversed the metabolic phenotype observed in mice lacking CnAβ1. Importantly, loss-of-function variants in CnAβ1 in humans were associated with similar metabolic changes. These findings suggest that targeting CnAβ1 expression could offer a novel therapeutic approach for combating obesity and other metabolic disorders, highlighting its potential as a key regulator in energy metabolism.

## RESULTS

### Lack of CnAβ1 accelerates brown pre-adipocyte differentiation

Since the ratio of CnAβ1 to CnAβ2 decreases during myoblast differentiation^15^, we investigated whether this was also the case in brown pre-adipocyte differentiation. We isolated pre-adipocytes from the BAT of wild type mice and differentiated them in culture. We found that the CnAβ1/CnAβ2 ratio decreased with differentiation (Fig. S1A). To gain further insight into the potential role of CnAβ1 in this process, we compared the differentiation capacity of pre-adipocytes isolated form wild type mice to those obtained from knockout mice lacking intron 12-13 in the *Ppp3cb* gene, which encodes the C-terminal domain in CnAβ1 (CnAβ1^Δi12^ mice; Fig. S1B-D). As shown in Fig. S1E, pre-adipocytes from CnAβ1^Δi12^ BAT differentiated faster than those from WT mice, as shown by the increased expression of Myf5, Pax7, Tbx20, Cox2, and Cpt1b mRNAs.

### Mice lacking CnAβ1 show reduced weight and smaller white adipocytes

To investigate the effect of the lack of CnAβ1 on whole body metabolism *in vivo*, we first bred WT and CnAβ1^Δi12^ mice on normal (ND) or high fat diet (HFD) and weighed them every 2 weeks after weaning until 12 weeks of age. CnAβ1^Δi12^ mice fed on HFD showed reduced weight compared to WT mice over time (Fig. 1A), while no differences were found in mice on ND at this age. We selected the final time point, 12 weeks old, to perform the rest of the experiments.

**Figure 1.**
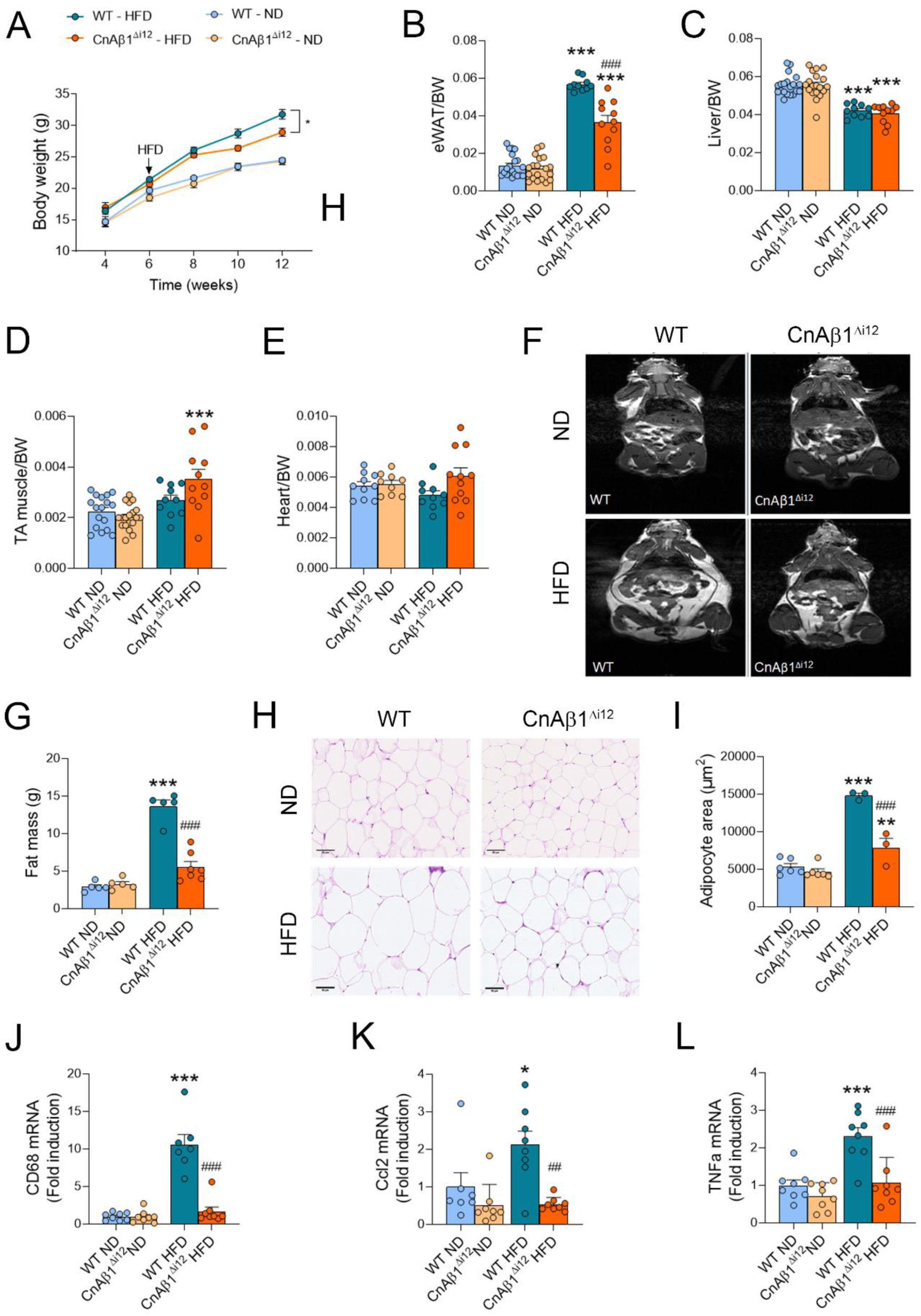
CnAβ1^Δi12^ mice show reduced body weight on HFD than WT mice and reduced white adipose tissue with smaller adipocytes. **A,** Mouse body weight was measured every two weeks from 4 to 12 weeks of age (n=7-16 per group) fed with normal (ND) or high fat diet (HFD). Results are expressed as mean ± SEM. *p<0.05 WT versus CnAβ1^Δi12^ mice, 2-way ANOVA plus Bonferroni post-test. **B-E**, Normalised weight of epidydimal white adipose tissue (eWAT; B), liver (C), tibialis anterior muscle (D), and heart (E) measured in CnAβ1^Δi12^ and WT mice on ND and HFD at 12 weeks of age (n=10-20 per group). **F, G,** Representative images (F) and quantification (G) of fat mass analysed by MRI. in WT and CnAβ1^Δi12^ mice (n=5-7). **H**, Representative H&E staining of eWAT isolated from mice fed with ND or HFD. Scale bar: 50 μm. **I**, Quantification of eWAT adipocyte cross sectional area mean (n=3-6 per group). **J-L**, CD68 (J), Ccl2 (K), and TNFα (L) mRNA expression was quantified by qRT-PCR. Results are expressed as mean ± SEM. *p<0.05, ***p<0.001 ND vs HFD; ^##^p<0.01, ^###^p<0.001 WT versus CnAβ1^Δi12^ mice, two-way ANOVA. eWAT: epidydimal white adipose tissue; TA, tibialis anterior.

To determine whether any particular tissue was responsible for the reduced weight in mice lacking CnAβ1, we measured the weight of various tissues that could influence total body weight. Epidydimal white fat pads in CnAβ1^Δi12^ mice on HFD showed a reduced weight compared to those of WT mice (Fig. 1B). In contrast, we did not find significant differences in the normalized weight of liver, tibialis anterior (TA) muscle and heart between CnAβ1^Δi12^ and WT mice at 12 weeks of age (Fig. 1C-E). Using MRI, we confirmed the smaller amounts of WAT in CnAβ1^Δi12^ (Fig. 1F, G). Histological analysis revealed a decreased size of white adipocytes in CnAβ1^Δi12^ mice (Fig. 1H, I), which also showed decreased expression of inflammatory markers in WAT (Fig. 1J-L). These results suggest that the difference in body weight between CnAβ1^Δi12^ and WT mice fed with HFD was due to a reduced white fat mass.

To explore potential physiological mechanisms that could explain the decreased weight of CnAβ1^Δi12^ mice on HFD, we housed the mice in metabolic cages for 60 hours. We observed a significant increase in O_2_ and CO_2_ exchange volumes and energy expenditure in CnAβ1^Δi12^ mice compared to WT mice on HFD (Fig. S2A-C). We found no significant changes in food intake or in physical activity in mice on HFD between both genotypes (Fig. S2D, E).

### CnAβ1^Δi12^ mice show decreased circulating glucose and triglycerides

To determine whether the absence of CnAβ1 prevented the deleterious effects of HFD on glucose uptake and/or insulin sensitivity, we performed GTT and ITT assays. We observed a trend of reduced blood glucose at every time point in both tests with a significant reduction in the area under the curve (AUC) in CnAβ1^Δi12^ mice when fed on a HFD (Fig. 2A-D). In addition, we found a decrease in blood glucose following overnight starvation (Fig. 2E), suggesting improved glucose uptake in agreement with the results observed in humans.

**Figure 2.**
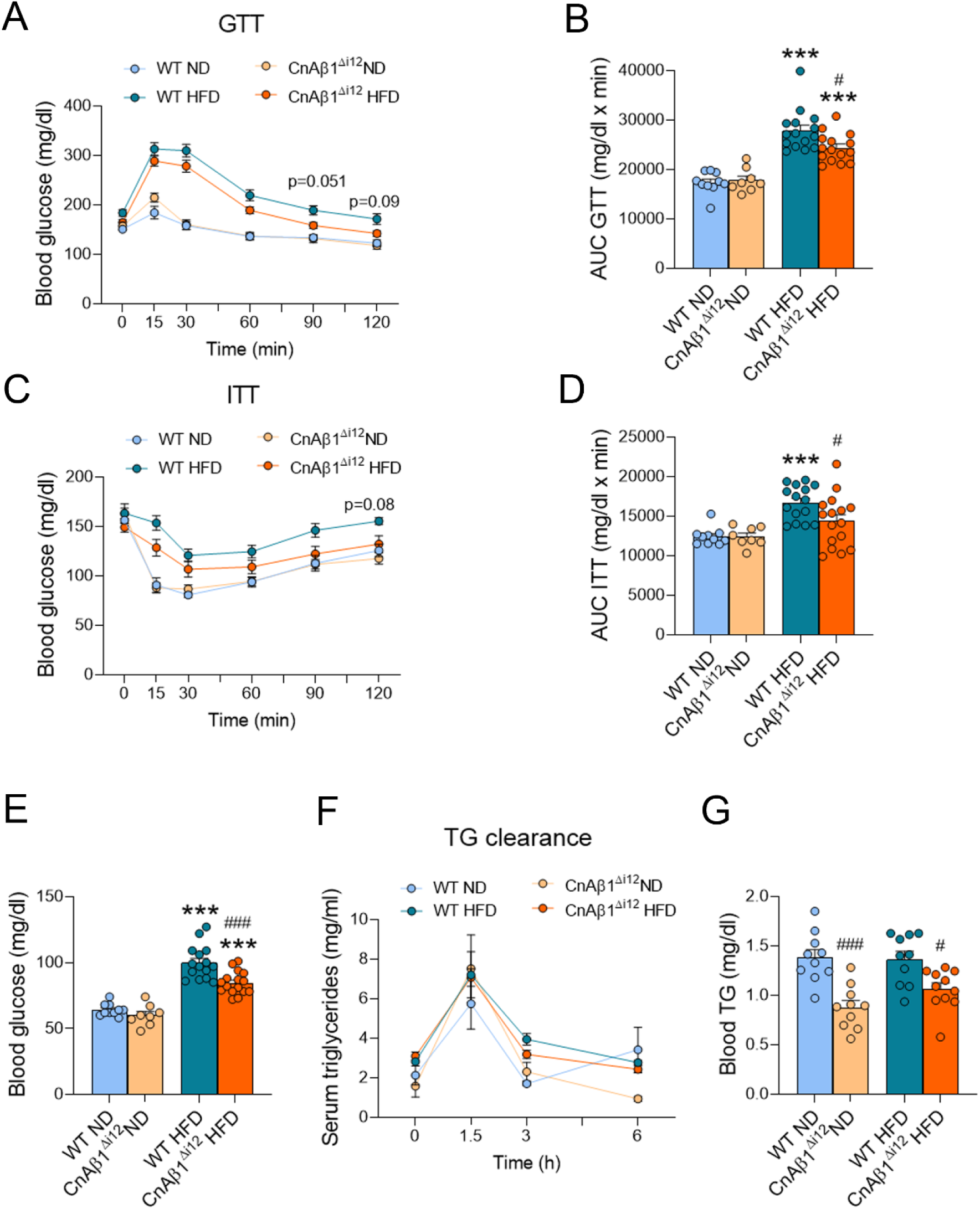
Glucose and fatty acid metabolism is improved in CnAβ1^Δi12^ mice fed with HFD. **A-D,** Glucose tolerance test (GTT, A, B) and insulin tolerance test (ITT, C, D) were performed in WT and CnAβ1^Δi12^ mice on ND and HFD. Mice were fasted for 5 h for GTT or 1 h for ITT, and 1 mg/g of glucose (GTT) or 0.75 UI/kg insulin (ITT) were injected i.p., respectively (n=7-15 per group). Blood glucose was measured at every time point indicated in the graph. C and D show the area under the curve (AUC) for each condition. **E**, Blood glucose was measured in WT and CnAβ1^Δi12^ mice (n=8-15). **F**, Serum TGs were measured at baseline (0) and 1.5, 3, and 6 h after administration by oral gavage of 15 ml/kg intralipid in overnight-fasted mice (n=8-10). **G**, Basal serum triglycerides were measured in WT and CnAβ1^Δi12^ mice (n=10). Results are expressed as mean ± SEM. ***p<0.001 ND vs HFD; ^#^p<0.05, ^###^p<0.001 WT versus CnAβ1^Δi12^ mice, two-way ANOVA plus Bonferroni post-test.

We next tested the capacity of white adipocytes to uptake triglycerides, which was similar in both genotypes (Fig. 2F), indicating that the decrease in white adipocyte size was not due to a defect or impairment in triglyceride uptake. However, CnAβ1^Δi12^ mice showed a reduction in blood triglycerides compared to WT mice, both on ND and HFD (Fig. 2G). These results pointed towards an accelerated usage of fatty acids elsewhere in the body.

### Increased mitochondria content and smaller lipid droplets in the BAT of CnAβ1^Δi12^ mice

Since the BAT is a major organ responsible for energy dissipation, we investigated whether it might be responsible for the improved metabolic phenotype of CnAβ1^Δi12^ mice on HFD. Mice lacking CnAβ1 showed increased normalized BAT weight on ND compared to WT mice, which was significantly decreased under HFD (Fig. 3A). Electron microscopy showed a decrease in the number of mitochondria in WT mice on HFD compared to ND that was prevented in the absence of CnAβ1 (Fig. 3B, 3C). CnAβ1^Δi12^ mice showed smaller lipid droplets than WT on HFD (Fig. 2D). Mitochondria were generally bigger in mice on HFD, with no major changes between both genotypes (Fig. 3E). Analysis of PGC1α, Lcad, TFAM, and Pnpla2/ATGL mRNA expression, associated with mitochondrial biogenesis, function and lipid catabolism, showed a reduction in WT mice on HFD, while it was preserved in CnAβ1^Δi12^ mice (Fig. 3F-H), thereby confirming the results observed by electron microscopy.

**Figure 3.**
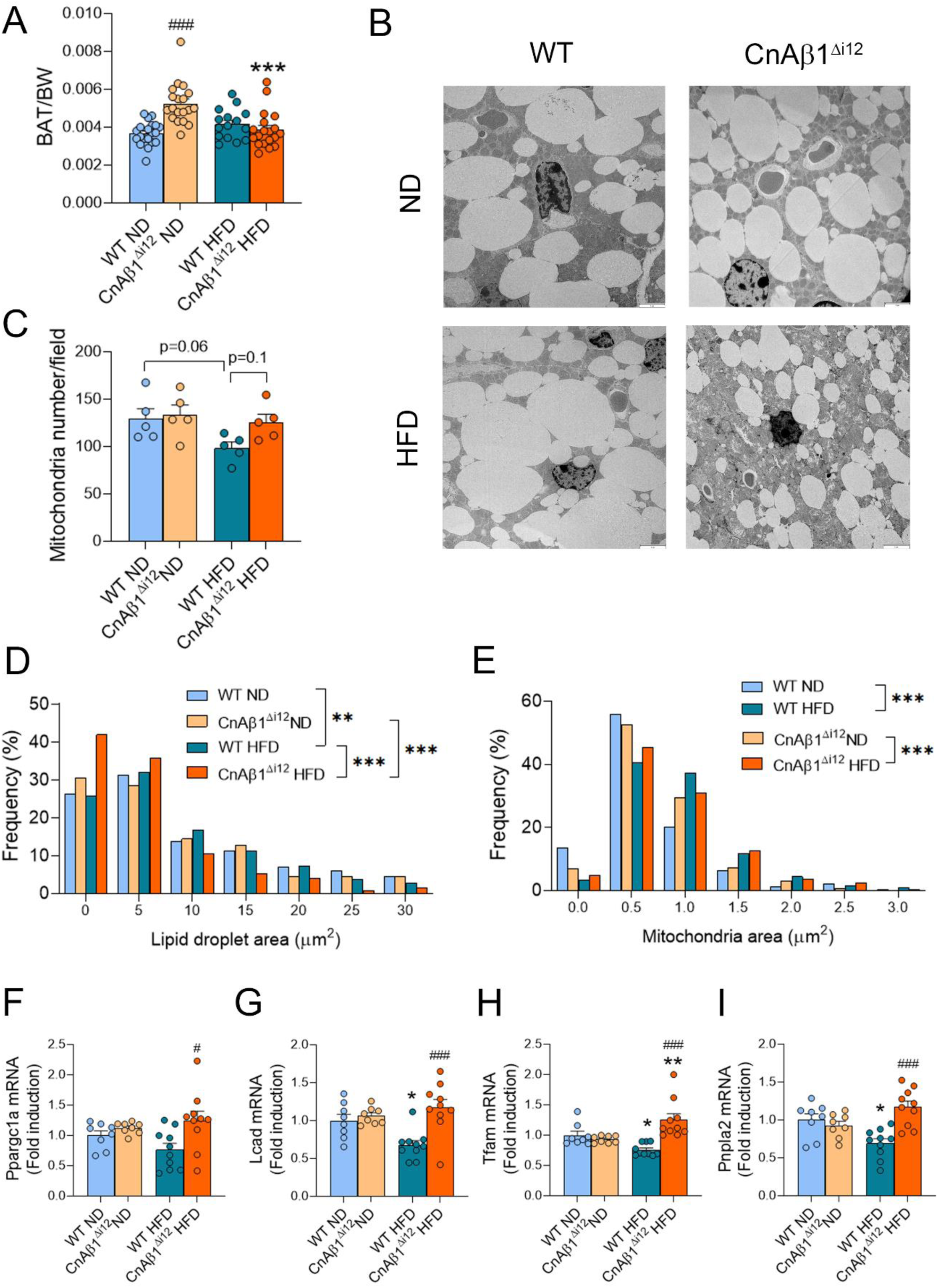
CnAβ1^Δi12^ mice show reduced lipid droplet size and more mitochondria in brown adipose tissue. **A**, BAT normalized weight measured in WT and CnAβ1^Δi12^ mice at 12 weeks of age after 6 weeks on ND or HFD (n=15-20). **B**, Electron microscope analysis of BAT from WT and CnAβ1^Δi12^ mice. Scale bar: 3 μm. **C-E**, The number of mitochondria (C), frequency of lipid droplet sizes (D), and frequency of mitochondrial areas were quantified in electron microscope images (n=5 per group). **F-I**, mRNA expression of Ppargc1a (encoding PGF1α; F), Lcad (G), Tfam (H), and Pnpla2 (encoding ATGL; I) in WT and CnAβ1^Δi12^ mice (n=8-10). Results are expressed as mean ± SEM. *p<0.05, **p<0.01, ***p<0.001 ND vs HFD; ^#^p<0.05, ^###^p<0.001 WT versus CnAβ1^Δi12^ mice, two-way ANOVA.

### Changes in the proteome of CnAβ1^Δi12^ mice are reminiscent of cold exposure and Rictor knockout

To gain insight into the mechanisms underlying the phenotypic changes in mice lacking CnAβ1, we analysed protein expression and post-translational modifications (PTMs) by mass-spectrometry proteomics in the BAT of WT and CnAβ1^Δi12^ mice fed with HFD. Loss of CnAβ1 resulted in an increased expression of proteins related to the mitochondria, fatty acid oxidation, and the electron transport chain, and thermogenesis (Table S1, Fig. 4A). Induction of the corresponding mRNAs was confirmed by qRT-PCR (Fig. S3A-C). CnAβ1^Δi12^ mice also showed increased expression of lipolysis genes (Pnpla2/Atgl, Lipe) and adipocyte differentiation markers (Cebpa, Pparg, Adipoq and Fabp4), suggesting that CnAβ1 regulated brown adipocyte differentiation, as suggested in the pre-adipocyte culture experiments.

**Figure 4.**
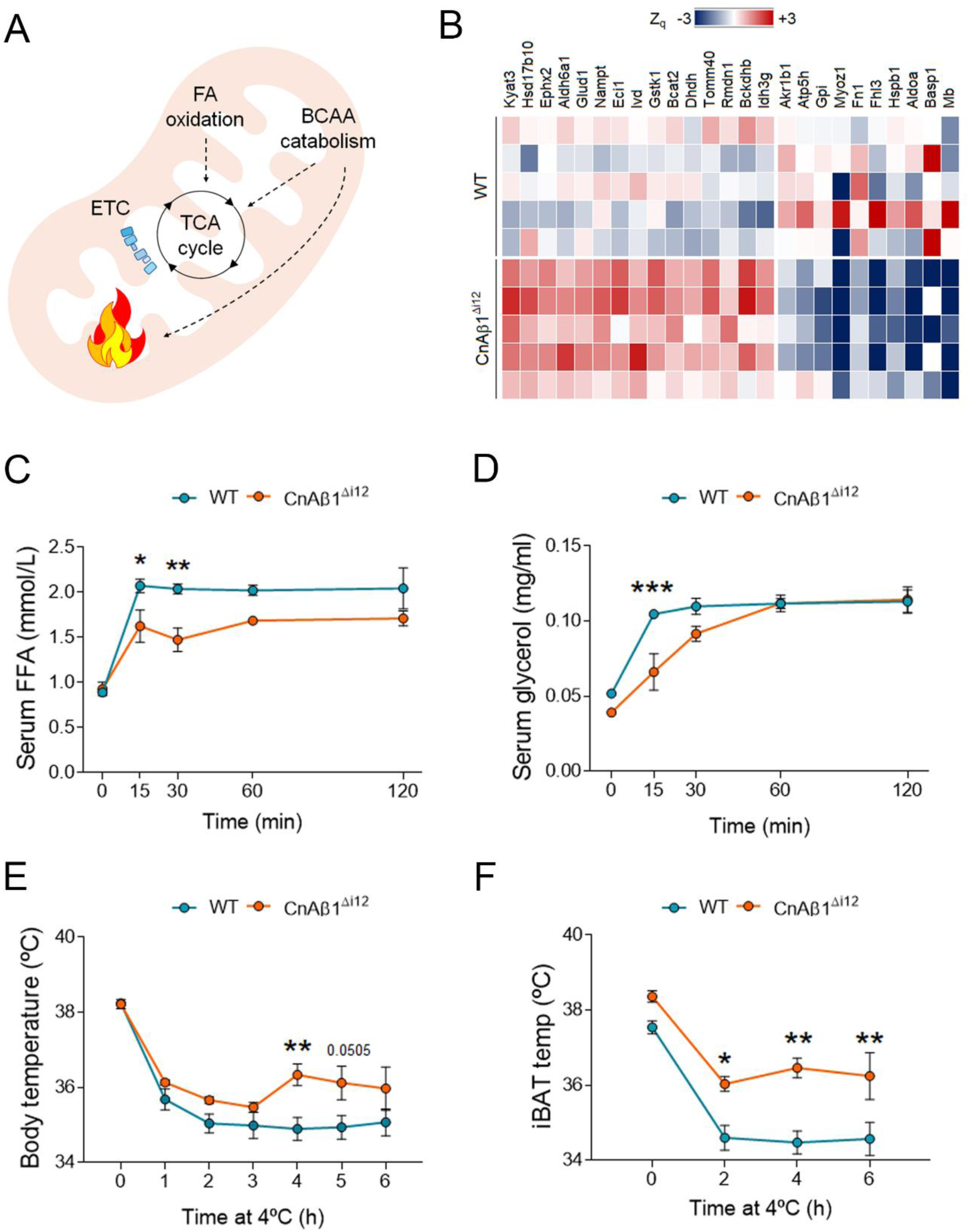
CnAβ1^Δi12^ mice on HFD show increased thermogenesis. **A**, Schematic summarising the upregulated proteins identified by proteomics in CnAβ1^Δi12^ mice. **B**, Top induced and reduced proteins in CnAβ1^Δi12^ mice compared to WT. **C, D**, WT and CnAβ1^Δi12^ mice on HFD (n=10 per group) were fasted for 3 h and intraperitoneal injections of β3 adrenergic agonist CL316.243 (1 mg/Kg) were administered. FFAs (C) and glycerol (D) were determined at the times indicated. Results are expressed as mean ± SEM. *p<0.05; **p<0.01; ***p<0.001 WT versus CnAβ1^Δi12^ mice, 2-way ANOVA plus Bonferroni post-test. **E, F**, Body (E) and interscapular BAT temperature (F) of WT and CnAβ1^Δi12^ mice (n=5 per group) after exposure to cold (4°C) for 6 h. Results are expressed as mean ± SEM. *p<0.05; **p<0.01 WT versus CnAβ1^Δi12^ mice, 2-way ANOVA plus Bonferroni post-test.

We investigated whether the increase in mitochondria-related genes observed in BAT was reproduced in other highly metabolic organs. We found no significant differences in the expression of these genes in liver between both mouse types (Fig. S4A). We found an increase in some mitochondria genes in skeletal muscle (2 out of 5), but no changes in thermogenic genes (Fig. S3B). No differences in mRNAs associated with different muscle fiber types or skeletal muscle differentiation were observed (Fig. S3C, S3D). These results suggest that the BAT is the main metabolic organ responsible for the changes observed in CnAβ1^Δi12^ mice.

In addition to genes involved in lipid catabolism, we observed in the proteomics analysis an upregulation of proteins associated with branched chain amino-acid (BCAA) catabolism (Table S1), which has been shown both to feed the TCA cycle to enhance thermogenesis and to improve glucose uptake in the liver in a thermogenic-independent manner^4,6^. The increase of the corresponding mRNAs was confirmed by qRT-PCR (Fig. S3D). We also observed enhanced expression of proteins associated with the production of kynurenic acid, which promotes energy expenditure and fatty acid oxidation in both BAT and WAT^17^. Down-regulated proteins included those associated with glycolysis and with skeletal muscle. Comparison of changes (Log_2_-fold change >0.6) in CnAβ1^Δi12^ vs WT mice with those induced by cold exposure vs room temperature^18^ revealed common induction of mitochondrial proteins and mitochondrial respiration (Table S1), and downregulation in both conditions of muscle-related genes (Table S1). This observed down-regulation of skeletal muscle proteins might be reminiscent of common Myf5^+^ precursor cells and has been described in response to cold exposure^18^. Interestingly, some of the changes observed in CnAβ1^Δi12^ mice were also observed in brown adipocytes from Rictor KO mice^19^ (Table S1).

Furthermore, we found that PLIN1 phosphorylation was inhibited in BAT of CnAβ1^Δi12^ mice (Table S2), suggesting post-translational regulation of lipolysis in this tissue. Reduced PLIN1 phosphorylation was also described in brown adipocytes from Rictor KO mice^19^. We also observed increased acetylation of some mitochondrial proteins in mice lacking CnAβ1 (Table S3), most of which were also more acetylated in the BAT of mice exposed to cold^18^. Since proteasome activity is necessary for BAT thermogenesis^20^, we next explored the changes in protein ubiquitination in CnAβ1^Δi12^ mice. We found increased ubiquitination in different mitochondrial proteins, including UCP1 (Table S4). These changes coincide with those previously observed after cold exposure^20^. In summary, the protein expression and post-translational modification changes observed in the absence of CnAβ1 largely reflect those described in mice exposed to cold.

### CnAβ1^Δi12^ mice are more resistant to cold

Stimulation of β3 adrenergic receptors leads to lipolysis in white adipose tissue and increased serum levels of FFA, which are used by the mitochondria in the BAT for heat production^21^. To investigate whether the lack of CnAβ1 had an impact on BAT activity, we stimulated mice with a β3 agonist. We observed a decrease in circulating FFAs and glycerol in CnAβ1^Δi12^ mice (Fig. 4B, 4C), suggesting that FFAs were being burnt by the BAT.

To study whether CnAβ1^Δi12^ mice produced more heat due to an increase in BAT temperature following β3 adrenergic stimulation, we exposed the mice to cold, which induces thermogenesis through the activation of β3 adrenergic receptors^22^. We housed the mice for 6 h at 4°C and we measured body and BAT temperature. As shown in Figure 5D and 4E, BAT temperature was increased in CnAβ1^Δi12^ mice compared to WT mice after a short-term cold exposure, accompanied by an increase in body temperature.

**Figure 5.**
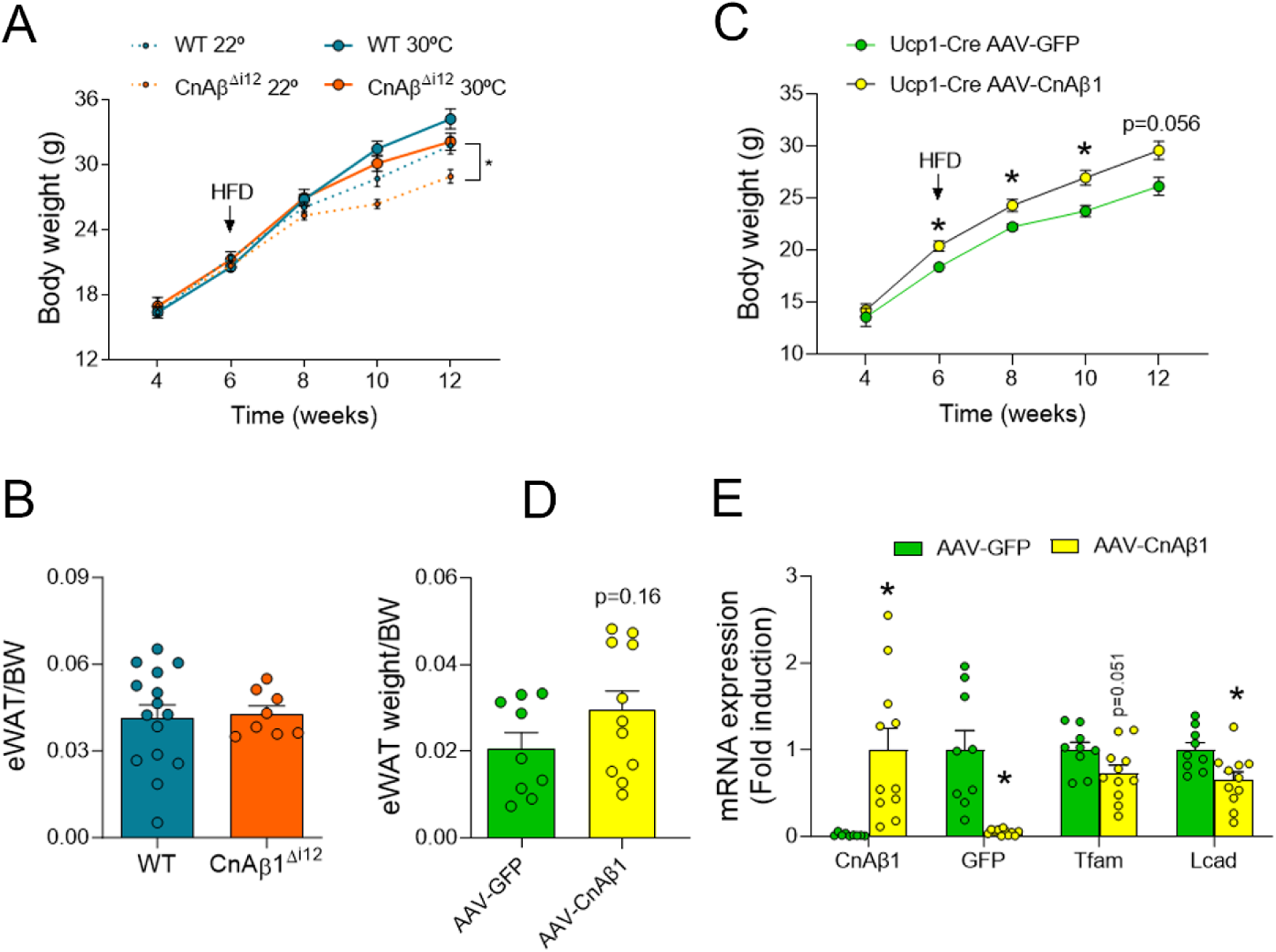
The BAT is partially responsible for the overall metabolic phenotype of CnAβ1^Δi12^ mice. **A**, Body weight was measured once every two weeks, starting at 4 weeks until 12 weeks of age in WT and CnAβ1^Δi12^ mice on HFD housed at 22°C or 30°C (n=8-15). Results are expressed as mean ± SEM. No significant differences were found between WT and CnAβ1^Δi12^ mice at 30°C, 2-way ANOVA plus Bonferroni post-test. **B**, Epididymal white adipose tissue weight to body weight ratio in WT (n=15) and CnAβ1^Δi12^ mice (n=8) on HFD housed at 30°C. Results are expressed as mean ± SEM. *p<0.05 WT versus CnAβ1^Δi12^ mice, unpaired Student’s t-test. **C**, CnAβ1^Δi12^ mice expressing the Cre recombinase under the control of the Ucp1 promoter (CnAβ1^Δi12^;Ucp1-Cre) were infected with AAV9-CnAβ1 or AAV9-GFP at 4 weeks of age. Mice were placed on HFD at 6 weeks and body weight was measured every 2 weeks (n=9-11). **D**, Epididymal white adipose tissue weight to body weight ratio in WT (n=9) and CnAβ1^Δi12^ mice (n=11) from the experiment in (C) measured at 12 weeks of age. **E**, mRNA expression of the indicated genes was analysed by qRT-PCR and normalized to β-actin mRNA in BAT at 12 weeks of age. *p<0.05 AAV9-CnAβ1 vs AAV9-GFP.

### The BAT is responsible for the reduced in body and WAT weight in CnAβ1^Δi12^ mice

Since CnAβ1^Δi12^ mice lack CnAβ1 expression in all tissues, it is not possible to determine whether a particular organ is responsible for the overall phenotype. To investigate whether the BAT was responsible for the weight changes observed in CnAβ1^Δi12^ mice, we first housed the mice at 30°C (thermoneutrality) starting after weaning, which is considered experimentally equivalent to the loss of BAT activity^23^. We found that the difference in body weight between WT and CnAβ1^Δi12^ mice at 12 weeks of age at 30°C (−2.08 g) had been reduced compared to the difference obtained at 22°C (−2.86 g; Fig. 5A). Similarly, the difference observed in eWAT/BW ratio between CnAβ1^Δi12^ and WT mice at 22°C (Fig. 1B) was lost when the mice were maintained in thermoneutral conditions (Fig. 5B).

Next, we investigated whether restoration of CnAβ1 expression specifically in the BAT of CnAβ1^Δi12^ mice would prevent the weight reduction in these mice. For this purpose, we crossed CnAβ1^Δi12^ mice with mice in which the Cre recombinase is expressed under the control of the BAT-specific Ucp1 promoter (Ucp1-Cre). We then infected the resulting CnAβ1^Δi12^;Ucp1-Cre mice with adeno-associated viral vectors (AAV9) harbouring either GFP or CnAβ1 cDNAs preceded by a floxed STOP codon, such that these proteins would only be expressed in tissues with an active Cre (i.e. those with an active Ucp1 promoter). Overexpression of CnAβ1 in BAT with AAV9-CnAβ1 resulted in a significant increase in body weight (Fig. 5C). In addition, restoration of CnAβ1 expression resulted in an increased eWAT/BW ratio and a reduction in at least some mitochondrial genes (Fig. 5D, 5E). These results suggest that the metabolic phenotype observed in CnAβ1^Δi12^ mice is at least partially due to the lack of CnAβ1 expression in BAT.

### FoxO1 mediates activation of the lipid catabolic programme in mice lacking CnAβ1

To determine the molecular mechanism underlying the changes in gene expression observed in CnAβ1^Δi12^ mice, we analysed different signalling pathways by western blot in BAT (Fig. 6A, 6B). We found a significant increase of PGC1α expression in CnAβ1^Δi12^ mice, confirming the qRT-PCR results. In addition, loss of CnAβ1 resulted in an increase in FoxO1 activation, shown as reduced phosphorylation and acetylation. This was paralleled by an inactivation of mTOR and Akt, as illustrated by reduced phosphorylation.

**Figure 6.**
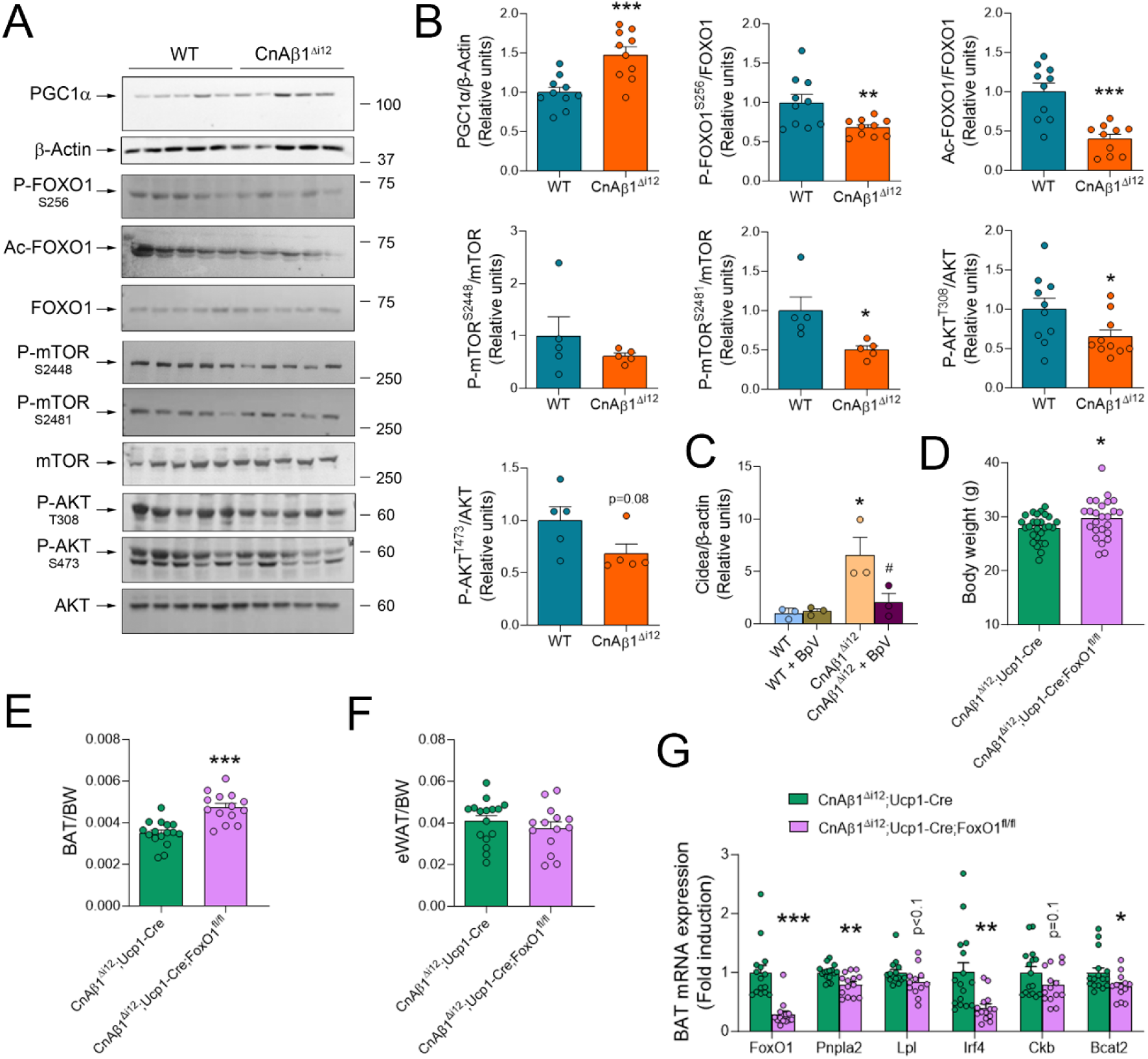
FoxO1 mediates the induction of lipid catabolism triggered by the absence of CnAβ1. **A, B**, Protein expression, phosphorylation and/or acetylation of the indicated proteins was analysed by western blot in WT and CnAβ1^Δi12^ mice after 6 weeks on HFD at 12 weeks of age. *p<0.05 WT vs CnAβ1^Δi12^, unpaired student’s t-test. **C**, Brown preadipocytes were differentiated towards adipocytes in the presence or absence of the PTEN inhibitor BpV. Cidea mRNA expression was analysed by qRT-PCR after differentiation. *p<0.05 WT vs CnAβ1^Δi12^; ^#^p<0.05 BpV vs control two-way ANOVA. n=3 per group. **D-G**, Role of FoxO1 in the catabolic phenotype induced by the absence of CnAβ1. Body weight (D), BAT (E) and eWAT (F) weight normalised to body weight in BAT-specific FoxO1 knockout CnAβ1^Δi12^ mice (CnAβ1^Δi12^;Ucp1-Cre;FoxO1^fl/fl^) compared to control mice (CnAβ1^Δi12^;Ucp1-Cre). BAT mRNA expression was analysed by qRT-PCR and normalized to β-actin mRNA at 12 weeks of age in CnAβ1^Δi12^;Ucp1-Cre;FoxO1^fl/fl^ compared to CnAβ1^Δi12^;Ucp1-Cre mice (G). *p<0.05, **p<0.01, ***p<0.001, unpaired t-test in CnAβ1^Δi12^;Ucp1-Cre;FoxO1^fl/fl^ compared to CnAβ1^Δi12^;Ucp1-Cre mice. n=14-16 per group.

To explore the functional role of the Akt/FoxO1 signalling axis in the changes observed in the BAT of CnAβ1^Δi12^ mice, we isolated BAT preadipocytes and differentiated them in culture. Adipocytes isolated from BAT in CnAβ1^Δi12^ mice showed significantly higher expression of the BAT differentiation marker Cidea compared to adipocytes form WT mice (Fig. 6C), confirming the changes observed in the BAT of in CnAβ1^Δi12^ mice. Adipocyte treatment with the PTEN inhibitor BpV, which activates the Akt pathway, prevented the increase in Cidea expression (Fig. 6C).

To further investigate the involvement of FoxO1 in the catabolic phenotype displayed by CnAβ1^Δi12^ mice, we crossed these mice with Ucp1-Cre;FoxO1^fl/fl^, which lack FoxO1 specifically in BAT. The resulting CnAβ1^Δi12^;Ucp1-Cre;FoxO1^fl/fl^ mice showed an increased body weight and a larger BAT compared to CnAβ1^Δi12^;Ucp1-Cre mice, while the eWAT remained unchanged (Fig. 6D-F).

We next looked for changes in catabolism-related genes genes and found decreased expression of the transcription factor Irf4 and genes involved in lipid and BCAA catabolism in CnAβ1^Δi12^;Ucp1-Cre;FoxO1^fl/fl^ mice (Fig. 6G). Together, these results suggest that FoxO1 activation in the BAT as a result of Akt inhibition is responsible for the activation of the lipid catabolic programme in the BAT of CnAβ1^Δi12^ mice.

### Reduced thermogenic capacity after fasting in mice lacking CnAβ1

Since CnAβ1^Δi12^ mice show an enhanced catabolic capacity, we investigated whether their response to fasting would be affected. We found that eWAT and BAT weight were both strongly reduced in CnAβ1^Δi12^ mice following overnight fasting, while the liver and TA muscle weight remained unaffected (Fig. 7A-D). Interestingly, CnAβ1^Δi12^ mice showed significantly reduced thermogenic capacity after fasting compared to WT mice (Fig. 7E-F).

**Figure 7.**
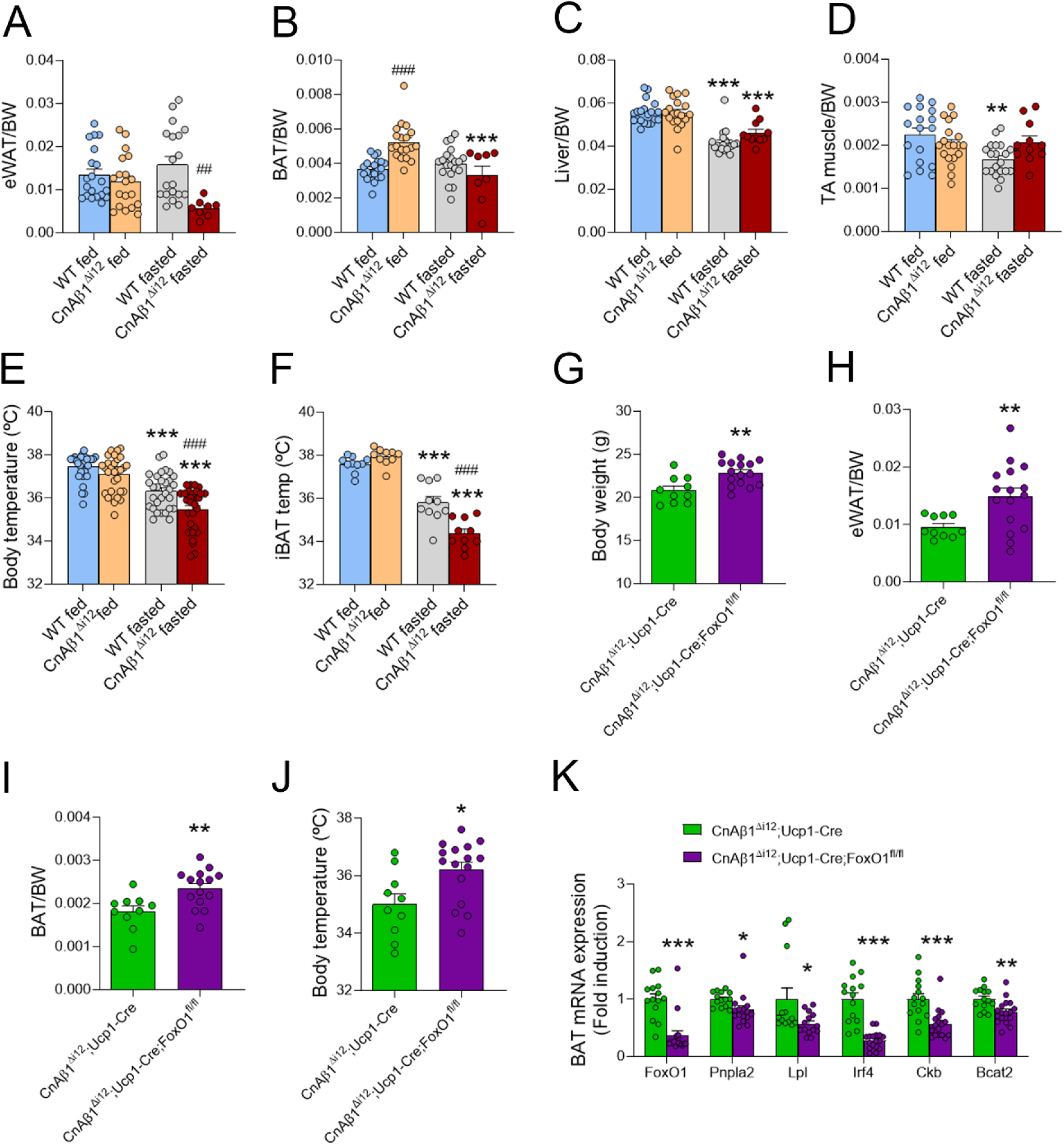
CnAβ1^Δi12^ mice lose body and BAT temperature after fasting. **A-D**, eWAT (A), BAT (B), liver (C) and tibialis anterior muscle (D) weight normalised to body weight was measured in WT and CnAβ1^Δi12^ mice fed with ND or after overnight fasting (n=11-20). **E, F**, Body (E; n=29-30) and intercapsular BAT temperature (F; n=10 per group) were measured in fed and fasted mice. **p<0.01, ***p<0.001 fed vs fasted; ^###^p<0.001 WT vs CnAβ1^Δi12^ mice. **G-I**, Body weight (G), eWAT (H), and BAT (I) weight normalised to body weight in BAT-specific FoxO1 knockout CnAβ1^Δi12^ mice (CnAβ1^Δi12^;Ucp1-Cre;FoxO1^fl/fl^) compared to control mice (CnAβ1^Δi12^;Ucp1-Cre) after overnight fasting. **J**, Body temperature in CnAβ1^Δi12^;Ucp1-Cre;FoxO1^fl/fl^ and control CnAβ1^Δi12^;Ucp1-Cre mice after overnight fasting. **K**, BAT mRNA expression was analysed by qRT-PCR and normalized to β-actin mRNA in CnAβ1^Δi12^;Ucp1-Cre;FoxO1^fl/fl^ compared to CnAβ1^Δi12^;Ucp1-Cre mice after overnight fasting. *p<0.05, **p<0.01, ***p<0.001, unpaired t-test in CnAβ1^Δi12^;Ucp1-Cre;FoxO1^fl/fl^) compared to CnAβ1^Δi12^;Ucp1-Cre mice. n=14-16 per group.

To determine whether BAT FoxO1 played a role in the accelerated lipid catabolism observed after fasting in CnAβ1^Δi12^ mice, we compared the response to overnight fasting of CnAβ1^Δi12^;Ucp1-Cre;FoxO1^fl/fl^ mice and CnAβ1^Δi12^;Ucp1-Cre controls. We found a significant increase in body weight and especially eWAT and BAT weight in CnAβ1^Δi12^;Ucp1-Cre;FoxO1^fl/fl^ mice (Fig. 7G-I). Furthermore, the absence of FoxO1 mitigated the fasting-induced hypothermia (Fig. 7J) and reduced the expression of genes involved in lipid and BCAA catabolism (Fig. 7K, L), suggesting that FoxO1 is mediating the overactivated catabolic programme triggered by the lack of CnAβ1.

### Loss-of-function variants in CnAβ1’s C-terminal domain are associated with improved metabolic function in humans

To explore the potential association between CnAβ1 and metabolic function in humans, we looked in UK Biobank for individuals with loss-of-function variants in the region of PPP3CB that encodes the C-terminal domain specific to CnAβ1. Our analysis revealed that these CnAβ1 loss-of-function variants were significantly associated with reduced blood glucose levels after correcting for age, sex and European ancestry (Fig. 8A). Additionally, we found a trend suggesting associations between these loss-of-function variants and lower BMI, reduced triglycerides, decreased cholesterol, and increased body temperature (Fig. 8B), although they were not statistically significant due to the limited number of subjects available. While further analysis was limited by the constraints of the available data, these trends confirmed the results found in mice, suggesting that CnAβ1 loss-of-function may have a beneficial effect on the regulation of body metabolism in humans.

**Figure 8.**
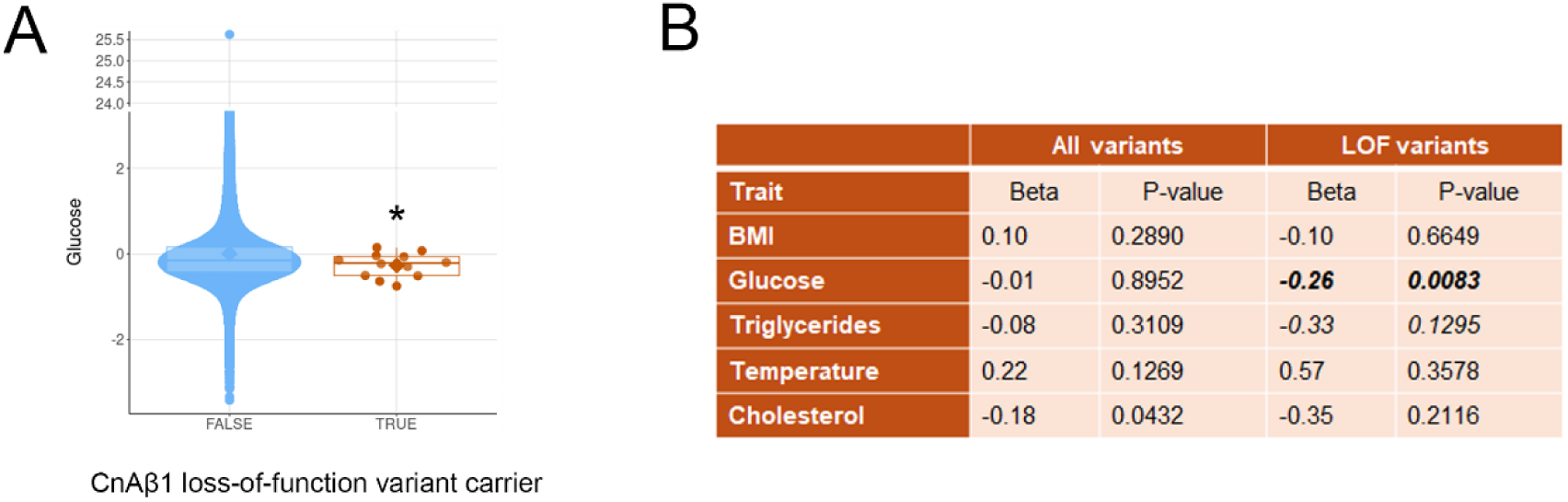
Loss-of-function variants in the C-terminal domain of CnAβ1 are associated with improved metabolic traits in humans. **A**, Blood glucose was significantly lower in human carriers of loss-of-function (LOF) variants in the C-ter domain of CnAβ1 (“TRUE”), compared to non-carriers (“FALSE”), in UK Biobank participants. **B**, Different metabolic parameters measured in carriers of LOF variants compared to non-carriers in UK Biobank participants.

## DISCUSSION

The role of calcineurin in adipose tissue metabolism is controversial. On one hand, mice deficient for the calcineurin inhibitor RCAN1, which have increased calcineurin activity, showed beigeing of WAT, reduced HFD-induced obesity, and improved insulin sensitivity^24^. In agreement with these results, a full knockout of the catalytic exon in the Ppp3cb gene, which affects the phosphatase activity of both CnAβ1 and CnAβ2, resulted in hyperlipidaemia and insulin resistance in mice on a regular chow diet^25^. Both reports therefore suggested that a loss of calcineurin activity is associated with metabolic dysregulation. Conversely, another study using the same Ppp3cb full knockout mice reported beneficial effects of reduced calcineurin activity when fed a HFD^26^, including improved insulin sensitivity. This study used a tissue-specific knockout of the calcineurin regulatory subunit CnB to exclude a direct role of calcineurin in lipolysis, thermogenesis, or adipogenesis in BAT or WAT, and instead suggested that the decrease in adipose tissue observed in Ppp3cb knockout mice was due to increased skeletal muscle activity^26^. No changes in BAT metabolism or in thermogenesis were reported. In addition, the tools used in these studies make it difficult to distinguish the role of each calcineurin isoform in adipose tissue metabolism, since 1) a CnB knockout affects all CnA subunits, 2) RCAN1 inhibits both CnAα and CnAβ2, and 3) the Ppp3cb exon 2 knockout would affect both CnAβ1 and CnAβ2. Furthermore, these studies failed to take into consideration the different CnAβ isoforms, which as mentioned above, have completely different mechanisms of action and which could account for the different phenotypes. This is potentially relevant, given the association of post-transcriptional regulation with different metabolic challenges recently unveiled^27^.

In this study, to differentiate the contribution of CnAβ1 from that of CnAβ2, we used mice lacking the intron encoding the C-terminal domain of CnAβ1. This approach allowed us to elucidate the previously unknown role of the alternative calcineurin variant CnAβ1 in regulating BAT metabolism. We found that the CnAβ1/CnAβ2 ratio decreases during brown pre-adipocyte differentiation. The absence of CnAβ1 accelerates pre-adipocyte differentiation and drives BAT overactivation, leading to a reduced body weight, smaller white adipocytes, improved glucose tolerance and enhanced cold resistance in mice fed with a HFD. Following starvation, this metabolic overactivation leads to a faster depletion of white fat depots and a reduced capacity to maintain body temperature. Importantly, we found similar trends in humans with loss-of-function variants in the C-terminal domain of CnAβ1.

Our findings suggest that the enhanced BAT activity and reduced body weight in CnAβ1^Δi12^ mice are at least partially due to the loss of CnAβ1 specifically in BAT. First, placing CnAβ1^Δi12^ mice in thermoneutral conditions mitigated the weight loss. Second, reintroducing CnAβ1 in BAT with a CnAβ1-overexpressing virus partially prevented the weight loss. Third, minimal changes in mitochondrial gene expression were observed in skeletal muscle and liver, suggesting that the phenotype is primarily driven by alterations in BAT. Finally, BAT adipocytes from CnAβ1^Δi12^ mice differentiated more effectively than those from wild type mice. While a direct effect of CnAβ1 on WAT cannot be entirely ruled out, our results strongly suggest that loss of CnAβ1 in BAT is the main contributor to the observed metabolic changes.

Unlike other calcineurin isoforms, CnAβ1 is palmitoylated and localized to cell membranes, where it interacts with mTOR^28,29^. CnAβ1 is necessary both for mTORC2 membrane localization and for the full activation of the Akt pathway^30,29^. Recent studies have shown that mTORC2 plays a critical role in regulating the expression of genes related to thermogenesis and mitochondrial function in BAT^31^. For example, mice with a BAT-specific deletion of the mTORC2 component Rictor showed increased expression of thermogenic genes, improved cold tolerance, and resistance to diet-induced obesity. mTORC2 inhibition also stimulated lipolysis and lipid uptake, thereby enhancing lipid catabolism and potentiating thermogenesis^31^. This effect was mediated by FOXO1 deacetylation and activation via the Sirt6 deacetylase^31^. Further studies from the same group found that deleting Rictor in Myf5^+^ cells, which are common precursors to both adipocytes and skeletal myocytes, increased diet-induced thermogenesis^32^. Similarly, PTEN transgenic mice, which have reduced Akt activity and increased FoxO1 activation in BAT, also showed activation of thermogenesis and a reduction in WAT size^33^. Consistent with these findings, we observed reduced Akt and enhanced FoxO1 activation in the BAT of CnAβ1^Δi12^ mice. BAT-specific FoxO1 knockout partially abolished the beneficial effects found in CnAβ1^Δi12^ mice fed a HFD, resulting in increased body weight. Similarly, loss of FoxO1 specifically in the BAT prevented the excessive energy depletion observed in CnAβ1^Δi12^ mice after starvation, restoring WAT depots and preventing hypothermia. In both conditions, the absence of FoxO1 in the BAT resulted in a decrease in genes involved in catabolic reactions. While we cannot exclude the activation of additional mechanisms, these results suggest that the reduced body weight in CnAβ1^Δi12^ mice was due to overactivation of the catabolic programme in the BAT as a result of FoxO1 activation.

Overall, this study unveils the role of CnAβ1 in regulating fat catabolism through the mTORC2-Akt-FoxO1 axis. The results highlight CnAβ1 as a novel potential target for reducing body weight and managing obesity and other metabolic disorders. By modulating CnAβ1 expression, it may be possible to enhance BAT activity and promote fat loss, offering a promising new avenue for therapeutic intervention.

## METHODS

### Mice

CnAβ1^Δi12^ mice were generated by removing intron 12-13 in the *CnAβ* gene, which encodes for the specific C-terminal domain of CnAβ1 (Figure S1A, S1B), and have been previously described^14^. These mice do not express CnAβ1 in fat tissues but a minimal increase in CnAβ2 expression is observed in epididymal white adipose tissue (eWAT) (Fig. S1C). Heterozygous animals were crossed with wild type C57BL/6J mice to obtain the tenth generation of CnAβ1^Δi12^ mice, and the resulting line was maintained in homozygosity. Adult male mice (12 weeks old) were used for all experiments to avoid the confounding effects of sex hormones and to reduce the number of mice. C57BL/6J male mice from Charles River (Sulzfeld, Germany) were used as controls. FoxO1 conditional knockout mice were derived from FoxO1/3/4 conditional knockout mice generated by Dr DePinho^34^. All procedures were approved by the local Ethics Committees of the Centro Nacional de Investigaciones Cardiovasculares (CNIC) and the Regional Government of Madrid (PROEX 177-17, PROEX191.5/22). Mice were housed in cages in standard laboratory conditions under a 12:12 h light/dark cycle, temperatures of 22°C ± 2°C, 40-60% relative humidity with free access to food and water. Animals were fed with normal (ND; Rod18-A LASQCdiet®) or high-fat diet (HFD; D12492i Research Diets), except when fasted for experiments. For all experiments, we fed the mice with HFD starting at 6 weeks until 12 weeks of age, making a total of 6 weeks on HFD. Body weight was measured every two weeks after the mice were weaned (4 weeks of age) until 12 weeks old.

### Magnetic Resonance Imaging (MRI)

Body fat mass was analysed by Magnetic Resonance Imaging performed by the Advance Imaging Unit of the CNIC. For image acquisition, animals were anesthetized with 2% isoflurane and 1.8 L/min oxygen flow. Mice eyes were covered with ophthalmic gel to prevent retinal drying. Mice were placed in a 7-T Agilent/Varian scanner (Agilent, Santa Clara, CA, USA) equipped with a DD2 console. Images were acquired by using an actively shielded 205/120 gradient with a volume transmit coil. Body fat imaging was obtained using a SEMS sequence with and without fat saturation with the following parameters: TR/TE = 533-884 /10.44 msec (TR varies between with and without fat saturation); acquisition matrix 128 x 128; flip angle 90°; 1 average; 35 slices; 1 mm slice thickness and slice distance. Image analysis was performed with Fiji software^35^. Briefly, each image representing a mouse was composed by 33 slices. The area of high intense pixels (fat) for each slice was calculated. This area was multiplied by the slice distance to obtain the fat volume, which was multiplied by 0.9 mg/ml (fat density) to obtain total value of fat mass in grams.

### Histology and electron microscopy

Tissues were isolated and fixed overnight in 4% paraformaldehyde/PBS (137 mM NaCl, 10 mM Na_2_HPO_4_-2H_2_O, 2.7 mM KCl, 1.8 mM KH_2_PO_4_). After fixation, tissues were submerged in 70% EtOH, dehydrated, embedded in paraffin, and sectioned in a Leica RM2135 microtome. 5 µm-thick tissue sections were processed for H&E staining by the Histology Unit at the CNIC. Adipocyte size and number were quantified using Fiji software on 20X magnification images. At least 20 adipocytes per mouse were selected to measure the cross-sectional area of white adipocytes. The average area of at least 60 adipocytes was calculated for each mouse. Histograms show the distribution of 60 adipocytes grouped by adipocyte cell area in each genotype.

For electron microscopy, sample were fixed in 1% glutaraldehyde, 4% formaldehyde in 0.1 M cacodylate buffer, and post-fixed in 1% osmium tetroxide in water. Samples were stained with 0.5% uranyl acetate in water, dehydrated progressively in 30%, 50%, 70%, 95%, 100% ethanol and acetone, and included in durcupan resin (Sigma-Aldrich). 60 nm-thick sections were obtained with a Leica Reichert Ultracut S ultramicrotome and counterstained with uranyl acetate and lead citrate. Sections were loaded on grids and examined using a Tecnai G2 spirit transmission electron microscope equipped with a lanthanum hexaboride (LaB6) filament and a TemCam-F416 (4k x 4k) camera with CMOS technology. The images were collected at nominal magnifications of 2410x and 9490x.

### Metabolic cages

Mice were individually housed in metabolic cages (OxyletPro-Physiocage from Panlab) for three days for acclimation before obtaining any measurement. Mice were then recorded for 60 h to measure O_2_ and CO_2_ exchange, energy expenditure, food intake and physical activity. All parameters were calculated and analyzed with Metabolism software (Panlab).

### Glucose and Insulin Tolerance Tests

Animals were fasted for 5 h before intraperitoneal injection of glucose (1 mg/g) for glucose tolerance test (GTT). Glucose was measured in blood samples obtained from the tail vein using a glucometer at baseline and at 15, 30, 60, 90 and 120 minutes after the glucose injection. For the insulin tolerance test (ITT), animals were fasted for 1 h before intraperitoneal injection of insulin (0.75 UI /Kg; Regular Humulin 100 UI/ml, Lilly). Serum glucose was measured in blood samples from the tail vein using a glucometer at baseline and at 15, 30, 60, 90 and 120 minutes after insulin injection.

### Triglyceride (TG) clearance and β_3_-adrenergic receptor stimulation

For the assessment of TG clearance, mice were overnight fasted before oral gavage of 3 mg/g of 20% intralipid (I141-100 ml; Sigma). Basal blood was extracted from the tail vein and at 1.5, 3 and 6 h after intralipid injection to measure serum triglycerides. Mice were fasted for 3 h before intraperitoneal injection of β3-agonist CL316.243 (1 mg/Kg; Sigma). Blood was extracted from the tail vein and analysed at baseline and at 15, 30, 60 and 120 minutes after β3-agonist injection to measure the levels of serum-free fatty acids and glycerol.

### Body and BAT temperature

Body temperature of mice was measured using a rectal probe (Ref. RET-3, World Precision Instruments) and interscapular BAT (iBAT) temperature was determined with a thermographic camera (T420, FLIR^®^). For this last procedure, animals were anesthetized with 2% isoflurane and the zone where the BAT was located was shaved. We took photographs of this zone with the thermographic camera. To obtain the average BAT temperature, ellipses were drawn in images around BAT and were analysed using FLIR® software.

### Cold and thermoneutrality exposure

For cold exposure, animals were individually housed in a fridge at 4°C for 6 h, and body and BAT temperature were measured every 1 or 2 hours, respectively. In thermoneutral conditions, animals were housed after weaning in a mouse rack set at 30°C until mice reached 12 weeks of age. Body weight was measured every 2 weeks since the animals were weaned. Final BAT and body temperature were also measured.

### Injection of adeno-associated viruses (AAV)

CnAβ1^Δi12^ mice were crossed with Ucp1-Cre mice for two generations to produce homozygous CnAβ1^Δi12^ mice expressing Cre under the control of the Ucp1 promoter. Male mice at 4 weeks of age were administered a retro-orbital injection of 50 μl containing 5×10^10^ vial particles of pAAVKEF1aHT-LSL-Egfp (control) or pAAVKEF1aHT-LSL-HA-CnAβ1 (overexpressing CnAβ1) under inhaled anaesthesia with isoflurane. These viruses have a STOP codon flanked by LoxP sites preceding the cDNA, such that the protein of interest is only expressed in the Cre-expressing tissue. Mice were fed with HFD starting at 6 weeks of age and sacrificed at 12 weeks of age.

### Determination of blood metabolites

Serum triglycerides and glycerol were measured with an enzymatic kit (TR0100; Sigma) following the manufacturer’s instructions. Triglycerides and glycerol standards were used from Sigma (Ref. G7793). Levels of free fatty acids were measured using a NEFA-HR assay test from FUJIFILM Wako Chemicals.

### Isolation, culture, and differentiation of preadipocytes from BAT

A total of 12 WT and 12 CnAβ1^Δi12^ male mice, 6 weeks old, were used. Brown adipose tissue was extracted from the interscapular region and, after cleaning and fixing with 100% methanol, it was treated with a 50:50 enzymatic preparation of isolation medium (175 ml PBS, 1M HEPES, P/S 100X, and fungizone 100X), and collagenase A solution (Collagenase from Clostridium histolyticum, C5138; Sigma Aldrich) at 15 mg/ml. The tissue was incubated in a 37°C water bath for 40 minutes, vortexed every 10 minutes, filtered through a 70 µm membrane (10788201, FalconTM 352350, Fisher Scientific), and centrifuged at 1500 g for 8 minutes at RT. The pellet was resuspended in maintenance medium (187.5 ml DMEM, 50 ml FBS 100%, 1M HEPES, P/S 100X, and fungizone 100X), and plated in a 12-well plate (734-2778, VWR). The plate was incubated at 37°C, with 5% CO_2_, 95% O_2_, and 95% relative humidity. After 7 days, differentiation was induced by adding differentiation medium (40 ml maintenance medium, 5 mg/ml insulin, and 20 μg/ml T3). Cells were incubated at 37°C for 96 hours, after which induction medium (differentiation medium, IBMX 500X, 3.9 mg/ml dexamethasone, and 44.75 mg/ml indomethacin) was added and cells were further incubated at 37°C for 72 hours. Where indicated, cells were incubated with 5 µM of the PTEN inhibitor BpV (HOpic; 203701, Sigma Aldrich) renewed with every medium change.

### RT-qPCR

RNA was extracted with TRIZOL (Invitrogen) using a TissueRuptor (Ref. 0003737000, IKA) and 100 ng of total RNA were used to synthesize the cDNA with the High-Capacity cDNA Reverse Transcription Kit (Ref. 4368814, Applied Biosystems). For cultured adipocytes, RNA was isolated using the Mini RNA Micro Kit (Ref. 217004, Qiagen). RT-qPCR was carried out in an Applied Biosystems 7900 Fast Real-Time PCR thermocycler using SybrGreen PCR Master Mix (Ref. 4309155, Applied Biosystems) and appropriate primers (Table 1). Changes in expression were calculated using the individual efficiency corrected calculated method using LinReg PCR Software^36^. Expression of each mRNA was normalized to β-actin and was expressed as relative fold-induction over the values of wild type mice.

### Western Blot

For protein extraction, tissue was homogenized with RIPA lysis buffer (50 mM Tris-HCl, 150 mM NaCl, 0.1% SDS, 1% NP40, 0.5% Deoxycholic Acid Sodium Salt, milliQ water) containing protease (Roche, ref. 05892791001) and phosphatase (Roche, ref. 04906837001) inhibitors. To help cell disruption and resuspension, Eppendorf micropestle and 1-hour tubes rotator at 4°C were used before tube centrifugation at maximum rpm at 4°C for 30 min to obtain lysates. Upper fat layer was removed and supernatant was used to measure protein quantity. Protein concentration was determined by Nanodrop and equal amounts of protein were separated in SDS-Acrylamide/Polyacrylamide gels using Tris-Glycine buffer. Proteins were transferred onto a PVDF membrane (IPVH00010, Millipore), blocked 1 h with 3% BSA or skimmed power milk in TBS 0.1% Tween-20 (TBST), incubated overnight at 4°C with the corresponding primary antibodies (Table 2), and incubated with HRP-conjugated secondary antibodies (Ref. P0447 for anti-mouse and Ref. P0448 for anti-rabbit, Dako) during 1 h at room temperature. Antibodies were diluted in TBST. Membranes were incubated for 1 min with ECL detection reagent (AC2204, Azure Biosystems). The IBright CL1500 imaging system (Thermofisher) was used to capture images and signal quantification was performed with Fiji software.

### Statistical analysis

GraphPad Prism 10 was used to perform all statistical analyses and graphs. After testing for normality with D’Agostino & Pearson test, unpaired Student’s t-test was used to compare means of two groups for single time point and two-way ANOVA with Bonferroni’s post-test was used to compare means of two groups for repeated measurements. For non-parametric data, Mann–Whitney U test was used as equivalent for unpaired Student’s t-test as indicated in figure legends. Values were represented as mean ± SEM and changes were considered significant when p-value <0.05.

## ACKNOWLEDGEMENTS

We are grateful to Jan-Bernd Funcke and Leon Straub for assistance with the generation of the AAV vectors. This work was partly supported by grants PID2021-124629OB-I00, PLEC2022-009235 funded by the Ministry of Science and Innovation (MCIN/ AEI/10.13039/501100011033), by the European Union’s NextGenerationEU/PRTR (“Plan de Recuperación, Transformación y Resiliencia de España”) and by Fondo Europeo de Desarrollo Regional (FEDER), and by grants PEJ-2023-TL/SAL-GL-28706 and PEJ-2019-TL/BMD-12831 from Comunidad de Madrid to E.L-P, and by Instituto de Salud Carlos III (PT23/00027 and PT20/00044) co-funded by European Union and European Regional Development Fund “A way to make Europe”. K.A.M. is supported by the British Heart Foundation [FS/IPBSRF/22/27059, RE/18/4/34215], and the NIHR Imperial College Biomedical Research Centre. J.S.W. is supported by the Sir Jules Thorn Charitable Trust [21JTA], Medical Research Council (UK) [MC_UP_1605/13], British Heart Foundation [RG/19/6/34387, RE/18/4/34215], and the NIHR Imperial College Biomedical Research Centre. This project used the ReDIB ICTS infrastructure TRIMA@CNIC, Ministerio de Ciencia e Innovación (MCIN). The CNIC is supported by the Instituto de Salud Carlos III (ISCIII), the Ministerio de Ciencia e Innovación (MCIN) and the Pro CNIC Foundation, and is a Severo Ochoa Center of Excellence (grant CEX2020-001041-S funded by MICIN/AEI/10.13039/501100011033).

## AUTHOR CONTRIBUTIONS

E.B., B.R., A.M., M.L-O., J.M.G-S., L.R., E.C., L.C., J.L.C., J.B., K.M., P.B. conceived and performed experiments, and carried out analyses. M.V.G-G, E.L.P. conceived experiments, wrote the manuscript, secured funding, administered the project, and supervised personnel. M.D., J.W., P.S., J.V., G.S. conceived experiments, supervised personnel, and secured funding.

## REFERENCES

1. Vegiopoulos, A., Rohm, M. & Herzig, S. Adipose tissue: between the extremes. EMBO J. 36, 1999–2017 (2017).

2. Henri, H. et al. Therapeutic Manipulation of Myocardial Metabolism. J. Am. Coll. Cardiol. 77, 2022–2039 (2021).

3. Kajimura, S. & Saito, M. A new era in brown adipose tissue biology: Molecular control of brown fat development and energy homeostasis. Annu. Rev. Physiol. 76, 225–249 (2014).

4. Yoneshiro, T. et al. BCAA catabolism in brown fat controls energy homeostasis through SLC25A44. Nature 572, 614–619 (2019).

5. Townsend, K. L. & Tseng, Y.-H. Brown fat fuel utilization and thermogenesis. Trends Endocrinol. Metab. 25, 168–177 (2014).

6. Verkerke, A. R. P. et al. BCAA-nitrogen flux in brown fat controls metabolic health independent of thermogenesis. Cell 187, 2359–2374.e18 (2024).

7. Becher, T. et al. Brown adipose tissue is associated with cardiometabolic health. Nat. Med. 27, 58–65 (2021).

8. Rusnak, F. & Mertz, P. Calcineurin: Form and function. Physiological Reviews vol. 80 1483–1521 at 10.1152/physrev.2000.80.4.1483 (2000).

9. Medyouf, H. & Ghysdael, J. The calcineurin/NFAT signaling pathway: A novel therapeutic target in leukemia and solid tumors. Cell Cycle vol. 7 297–303 at 10.4161/cc.7.3.5357 (2008).

10. Gomez-Salinero, J. M., García-Pavía, P. & Lara-Pezzi, E. CnAβ1 shifts cardiac metabolism. Aging (Albany. NY). 11, 839–840 (2019).

11. Lara-Pezzi, E. et al. A naturally occurring calcineurin variant inhibits FoxO activity and enhances skeletal muscle regeneration. J. Cell Biol. 179, 1205–1218 (2007).

12. López-Olañeta, M. M. et al. Induction of the calcineurin variant CnAβ1 after myocardial infarction reduces post-infarction ventricular remodelling by promoting infarct vascularization. Cardiovasc. Res. 102, 396–406 (2014).

13. Felkin, L. E. et al. Calcineurin splicing variant calcineurin Aβ1 improves cardiac function after myocardial infarction without inducing hypertrophy. Circulation 123, 2838–47 (2011).

14. Padrón-Barthe, L. et al. Activation of serine one-carbon metabolism by calcineurin Aβ1 reduces myocardial hypertrophy and improves ventricular function. J. Am. Coll. Cardiol. 71, 654–667 (2018).

15. Lara-Pezzi, E. et al. A naturally occurring calcineurin variant inhibits FoxO activity and enhances skeletal muscle regeneration. J. Cell Biol. 179, 1205–1218 (2007).

16. Seale, P. et al. PRDM16 controls a brown fat/skeletal muscle switch. Nature 454, 961–967 (2008).

17. Agudelo, L. Z. et al. Kynurenic Acid and Gpr35 Regulate Adipose Tissue Energy Homeostasis and Inflammation. Cell Metab. 27, 378–392.e5 (2018).

18. Entwisle, S. W., et al. Cold-Induced Thermogenesis Increases Acetylation on the Brown Fat Proteome and Metabolome. bioRxiv 445718 (2018) doi:10.1101/445718.

19. Entwisle, S. W. et al. Proteome and Phosphoproteome Analysis of Brown Adipocytes Reveals That RICTOR Loss Dampens Global Insulin/AKT Signaling*. Mol. Cell. Proteomics 19, 1104–1119 (2020).

20. Bartelt, A. et al. Brown adipose tissue thermogenic adaptation requires Nrf1-mediated proteasomal activity. Nat. Med. 24, 292–303 (2018).

21. Festuccia, W. T., Blanchard, P.-G. & Deshaies, Y. Control of Brown Adipose Tissue Glucose and Lipid Metabolism by PPARγ. Front. Endocrinol. (Lausanne). 2, 84 (2011).

22. Lowell, B. B. & Spiegelman, B. M. Towards a molecular understanding of adaptive thermogenesis. Nature vol. 404 652–660 at 10.1038/35007527 (2000).

23. Cui, X. et al. Thermoneutrality decreases thermogenic program and promotes adiposity in high-fat diet-fed mice. Physiol. Rep. 4, 12799 (2016).

24. Rotter, D. et al. Regulator of Calcineurin 1 helps coordinate whole-body metabolism and thermogenesis. EMBO Rep. 19, e44706 (2018).

25. Suk, H. Y. et al. Ablation of calcineurin Aβ reveals hyperlipidemia and signaling cross-talks with phosphodiesterases. J. Biol. Chem. 288, 3477–3488 (2013).

26. Pfluger, P. T. et al. Calcineurin links mitochondrial elongation with energy metabolism. Cell Metab. 22, 838–850 (2015).

27. Li, J. et al. Isoform usage as a distinct regulatory layer driving nutrient-responsive metabolic adaptation. Cell Metab. 37, 772–787.e6 (2025).

28. Ulengin-Talkish, I. et al. Palmitoylation targets the calcineurin phosphatase to the phosphatidylinositol 4-kinase complex at the plasma membrane. Nat. Commun. 12, 6064 (2021).

29. Gómez-Salinero, J. M. et al. The Calcineurin Variant CnAβ1 Controls Mouse Embryonic Stem Cell Differentiation by Directing mTORC2 Membrane Localization and Activation. Cell Chem. Biol. 23, 1372–1382 (2016).

30. Lara-Pezzi, E. et al. A naturally occurring calcineurin variant inhibits FoxO activity and enhances skeletal muscle regeneration. J. Cell Biol. 179, 1205–1218 (2007).

31. Jung, S. M. et al. Non-canonical mTORC2 Signaling Regulates Brown Adipocyte Lipid Catabolism through SIRT6-FoxO1. Mol. Cell 75, 807–822.e8 (2019).

32. Hung, C. M. et al. Rictor/mTORC2 loss in the Myf5 lineage reprograms brown fat metabolism and protects mice against obesity and metabolic disease. Cell Rep. 8, 256–271 (2014).

33. Ortega-Molina, A. et al. Pten Positively Regulates Brown Adipose Function, Energy Expenditure, and Longevity. Cell Metab. 15, 382–394 (2012).

34. Paik, J.-H. et al. FoxOs Are Lineage-Restricted Redundant Tumor Suppressors and Regulate Endothelial Cell Homeostasis. Cell 128, 309–323 (2007).

35. Schindelin, J., et al. Fiji: an open-source platform for biological-image analysis. Nat. Methods 9, 676–682 (2012).

36. Rao, X., Huang, X., Zhou, Z. & Lin, X. An improvement of the 2^(−delta delta CT) method for quantitative real-time polymerase chain reaction data analysis. Biostat. Bioinforma. Biomath. 3, 71–85 (2013).

