## Supplementary Information for "Loss of the alternative calcineurin variant CnAβ1 enhances brown adipocyte differentiation and drives metabolic overactivation through FoxO1 activation"

**ONLINE SUPPLEMENT**

### SUPPLEMENTARY METHODS

*Association of CnAβ1 genetic variants with metabolic parameters*

The UK Biobank cohort – The UK Biobank (UKB) recruited 500,000 participants aged 40–69 years across the UK between 2006 and 2010 (National Research Ethics Service, 11/NW/0382, 21/NW/0157). Written informed consent was provided. This study was conducted under terms of access approval number 47602. 469,835 UKB participants underwent exome sequencing[1].

Exome sequencing data analysis and variant curation – Variants within the region of GrCh38 73,439,902-73,444,724 on chromosome 10 were extracted from the exome sequencing data. All variants were covered by the sequencing, the maximum missing values from quality control was 0.28%. Protein-altering variants in the region 73,444,600-73,444,724, which codes for the C-terminal domain in CnAβ1, that had a MAF of <0.1% in gnomAD and UKB were identified. All variants passed UKB quality control (ukb23158_500k_OQFE.90pct10dp_qc_variants.txt). The exome sequencing data was annotated using Ensembl Variant Effect Predictor (VEP; version 104) with a plugin for gnomAD (version r2.1), and the data was organised using PLINK (version 1.90p 64-bit). The VEP output was analysed using R (version 3.6.0) and Rstudio (version 1.3.1073).

UKB codes for phenotype analysis – Measures of glucose (UK Biobank trait ID 30740-0.0) at recruitment, triglycerides (30870-0.0) at recruitment, body mass index (BMI; 21001-0.0) at recruitment, and average temperature (90192-0.0) on accelerometer, were analysed from the UK Biobank data. Glucose and triglycerides were analysed by blood biochemistry and measured by hexokinase analysis and GPO-POD analysis, respectively, on a Beckman Coulter AU5800. The BMI value is constructed from height and weight measured during the initial Assessment Centre visit. Temperature of device from the physical activity measurement recorded via a wrist-worn accelerometer[2] was assessed.

Statistical analysis – The association between variant carrier status and phenotypes were tested using Student's *t*-test. The phenotypes were adjusted for age at recruitment (21022-0.0), sex (22001-0.0), and genetic-derived European ancestry[3] in a linear fashion with residuals standardised as mean=0 and SD=1. All analyses were undertaken using R software. Two-sided student's t-test was used to test for a difference in means and Fischer exact test was used to assess categorical variables. P-values were adjusted by Bonferroni correction for the four phenotypes tested.

*Proteomics analyses*

Proteins were extracted from brown adipose tissue samples as previously described [4], and protein concentration in the resulting preparations was determined using the RCDC Protein Assay Kit (Bio-Rad, Hercules, CA, USA). Protein tryptic digestion was carried out using 30 K FASP filters (Abcam, Cambridge, UK) following the manufacturer's instructions [5], after which the resulting peptides were isobarically labeled with 18-plex Tandem Mass Tags (TMT, Thermo Scientific, Waltham, MA USA) reagents following the manufacturer's instructions [6]. An aliquot of the labeled peptides was separated into five fractions using the high pH fractionation kit (Thermo Scientific) [7]. Liquid chromatography tandem mass spectrometry (LC-MS/MS) analysis was performed on an Easy-nLC 1000 HPLC system (Thermo Fisher Scientific) coupled via a nanoelectrospray ion source (Thermo Fisher Scientific) to a Q Exactive HF mass spectrometer (Thermo Fisher Scientific) as described elsewhere[8], and the raw LC-MS/MS data were matched against a Uniprot Mus musculus database (January 2022; 65,300 entries) with Proteome Discoverer (version 2.5, Thermo Fisher Scientific) for peptide identification[9,10] including the following variable modifications: Met oxidation; phosphorylation at Ser, Thr, and Tyr; Lys acetylation; and Lys ubiquitination. Then the statistical assessment of protein and peptide abundance changes was carried out using the iSanXoT software package [11-13]. Finally, the Limma package [14] was used to ascertain statistical significance by means of p-values.

### REFERENCES


1. Backman, J.D., Li, A.H., Marcketta, A., Sun, D., Mbatchou, J., Kessler, M.D., Benner, C., Liu, D., Locke, A.E., Balasubramanian, S., et al. (2021). Exome sequencing and analysis of 454,787 UK Biobank participants. Nature *599*, 628-634. 10.1038/s41586-021-04103-z.

2. van Hees, V.T., Fang, Z., Langford, J., Assah, F., Mohammad, A., da Silva, I.C.M., Trenell, M.I., White, T., Wareham, N.J., and Brage, S. (2014). Autocalibration of accelerometer data for free-living physical activity assessment using local gravity and temperature: an evaluation on four continents. Journal of Applied Physiology *117*, 738-744. 10.1152/japplphysiol.00421.2014.

3. Meyer, H.V., Dawes, T.J.W., Serrani, M., Bai, W., Tokarczuk, P., Cai, J., de Marvao, A., Henry, A., Lumbers, R.T., Gierten, J., et al. (2020). Genetic and functional insights into the fractal structure of the heart. Nature *584*, 589-594. 10.1038/s41586-020-2635-8.

4. Diaz Marin, R., Crespo-Garcia, S., Wilson, A.M., and Sapieha, P. (2019). RELi protocol: Optimization for protein extraction from white, brown and beige adipose tissues. MethodsX *6*, 918-928. 10.1016/j.mex.2019.04.010.

5. Wisniewski, J.R. (2018). Filter-Aided Sample Preparation for Proteome Analysis. Methods Mol Biol *1841*, 3-10. 10.1007/978-1-4939-8695-8_1.

6. Li, J., Van Vranken, J.G., Pontano Vaites, L., Schweppe, D.K., Huttlin, E.L., Etienne, C., Nandhikonda, P., Viner, R., Robitaille, A.M., Thompson, A.H., et al. (2020). TMTpro reagents: a set of isobaric labeling mass tags enables simultaneous proteome-wide measurements across 16 samples. Nat Methods *17*, 399-404. 10.1038/s41592-020-0781-4.

7. Wang, Y., Yang, F., Gritsenko, M.A., Wang, Y., Clauss, T., Liu, T., Shen, Y., Monroe, M.E., Lopez-Ferrer, D., Reno, T., et al. (2011). Reversed-phase chromatography with multiple fraction concatenation strategy for proteome profiling of human MCF10A cells. Proteomics *11*, 2019-2026. 10.1002/pmic.201000722.

8. Binek, A., Castans, C., Jorge, I., Bagwan, N., Rodriguez, J.M., Fernandez-Jimenez, R., Galan-Arriola, C., Oliver, E., Gomez, M., Clemente-Moragon, A., et al. (2024). Oxidative Post-translational Protein Modifications upon Ischemia/Reperfusion Injury. Antioxidants (Basel) *13*. 10.3390/antiox13010106.

9. Bonzon-Kulichenko, E., Garcia-Marques, F., Trevisan-Herraz, M., and Vazquez, J. (2015). Revisiting peptide identification by high-accuracy mass spectrometry: problems associated with the use of narrow mass precursor windows. Journal of proteome research *14*, 700-710. 10.1021/pr5007284.

10. Navarro, P., and Vazquez, J. (2009). A refined method to calculate false discovery rates for peptide identification using decoy databases. Journal of proteome research *8*, 1792-1796. 10.1021/pr800362h.

11. Garcia-Marques, F., Trevisan-Herraz, M., Martinez-Martinez, S., Camafeita, E., Jorge, I., Lopez, J.A., Mendez-Barbero, N., Mendez-Ferrer, S., Del Pozo, M.A., Ibanez,

B., et al. (2016). A Novel Systems-Biology Algorithm for the Analysis of Coordinated Protein Responses Using Quantitative Proteomics. Mol Cell Proteomics *15*, 1740-1760. 10.1074/mcp.M115.055905.

12. Navarro, P., Trevisan-Herraz, M., Bonzon-Kulichenko, E., Nunez, E., Martinez-Acedo, P., Perez-Hernandez, D., Jorge, I., Mesa, R., Calvo, E., Carrascal, M., et al. (2014). General statistical framework for quantitative proteomics by stable isotope labeling. Journal of proteome research *13*, 1234-1247. 10.1021/pr4006958.

13. Rodriguez, J.M., Jorge, I., Martinez-Val, A., Barrero-Rodriguez, R., Magni, R., Nunez, E., Laguillo, A., Devesa, C.A., Lopez, J.A., Camafeita, E., and Vazquez, J. (2024). iSanXoT: A standalone application for the integrative analysis of mass spectrometry-based quantitative proteomics data. Comput Struct Biotechnol J *23*, 452-459. 10.1016/j.csbj.2023.12.034.

14. Ritchie, M.E., Phipson, B., Wu, D., Hu, Y., Law, C.W., Shi, W., and Smyth, G.K. (2015). limma powers differential expression analyses for RNA-sequencing and microarray studies. Nucleic Acids Res *43*, e47. 10.1093/nar/gkv007.

### SUPPLEMENTARY FIGURES

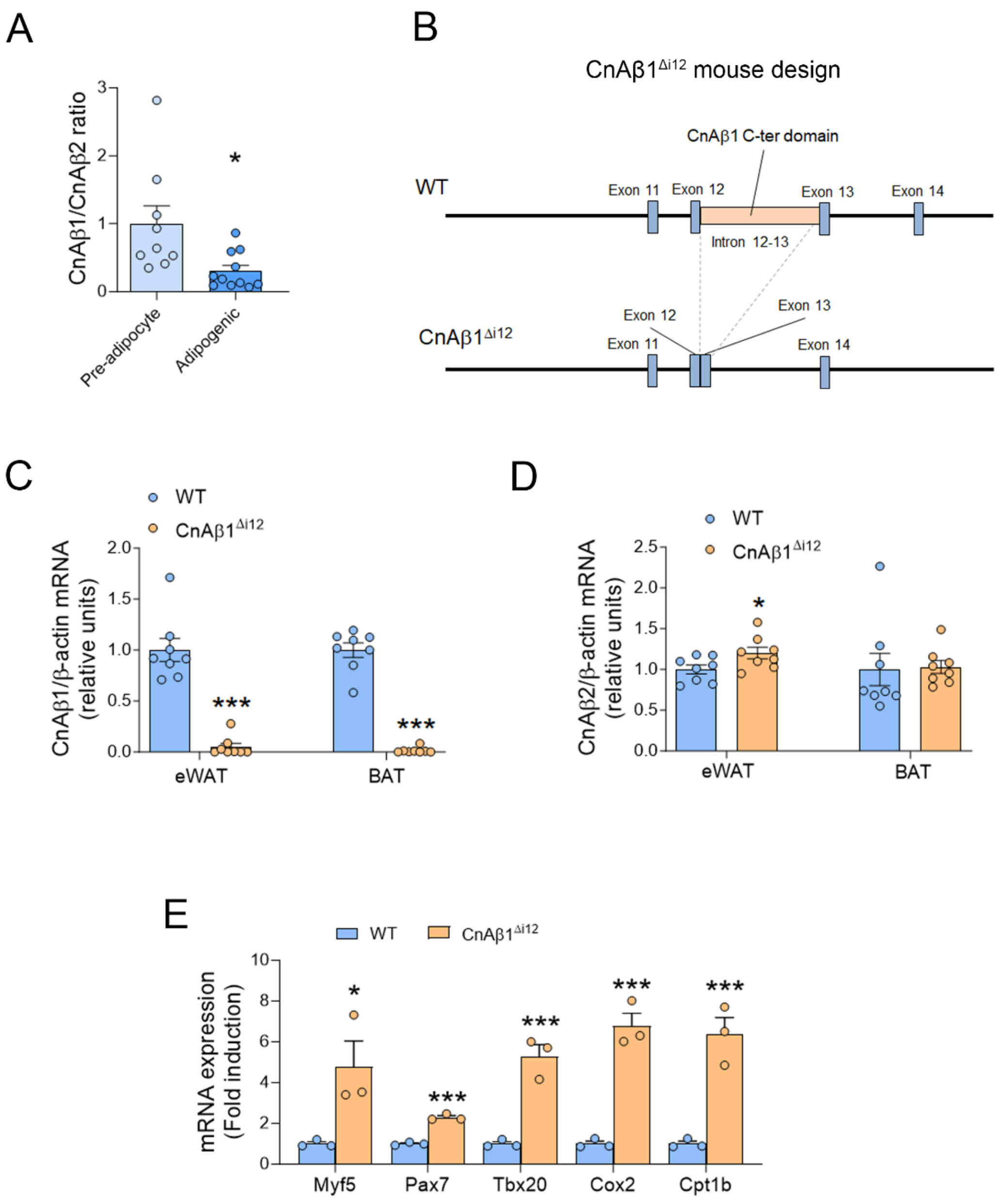


**Figure S1. Accelerated differentiation in brown pre-adipocytes from CnAβ1$^{\Delta 12}$ mice. A**, Blood glucose was significantly lower in human carriers of loss-of-function variants in the C-ter domain of CnAβ1. B, Schematic showing how CnA□1□i12 mice were developed by removing intron 12-13 in CnAβ. C, D Expression of CnAβ1 (B) and CnAβ2 (C) mRNA was determined by qRT-PCR in eWAT and BAT and normalise to that of β-Actin. E-G, Liver (D), tibialis anterior (E), and heart (E) weight normalised to body weight in wild type and CnAβ1$^{\Delta i12}$ mice fed a normal or high-fat diet. *p<0.05, ***p<0.001 normal diet vs high-fat diet.

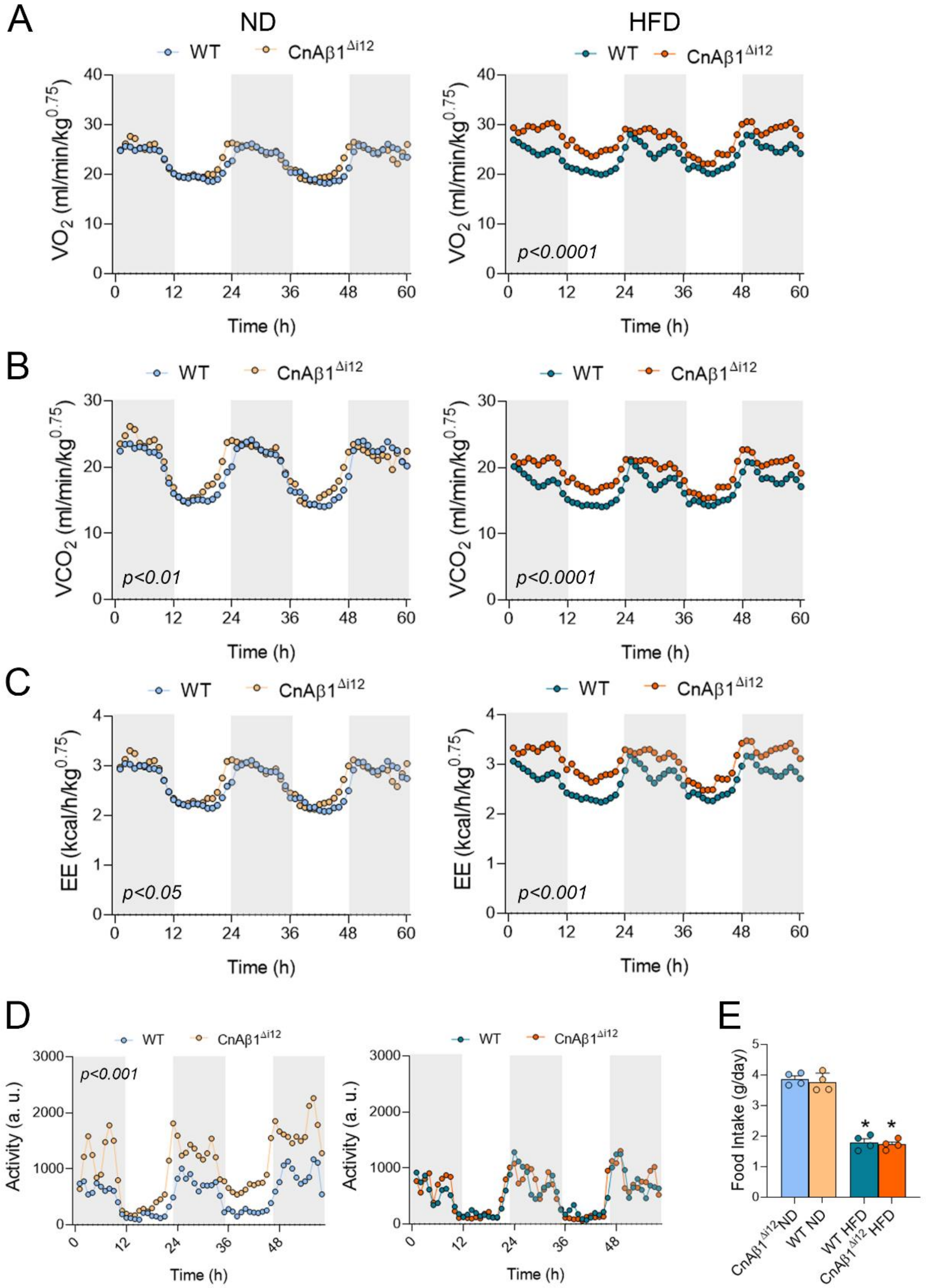


**Figure S2. CnAβ1$^{\Delta i12}$ mice fed a HFD show a mild increase in energy expenditure compared to WT mice. A-E**, Metabolic cages were used to quantify the volume of $O_2$ exchange (A), $CO_2$ exchange (B), energy expenditure (C), physical activity (D) and food intake (E). The p-values refer to WT vs CnAβ1$^{\Delta i12}$ mice using a 2-way ANOVA test.

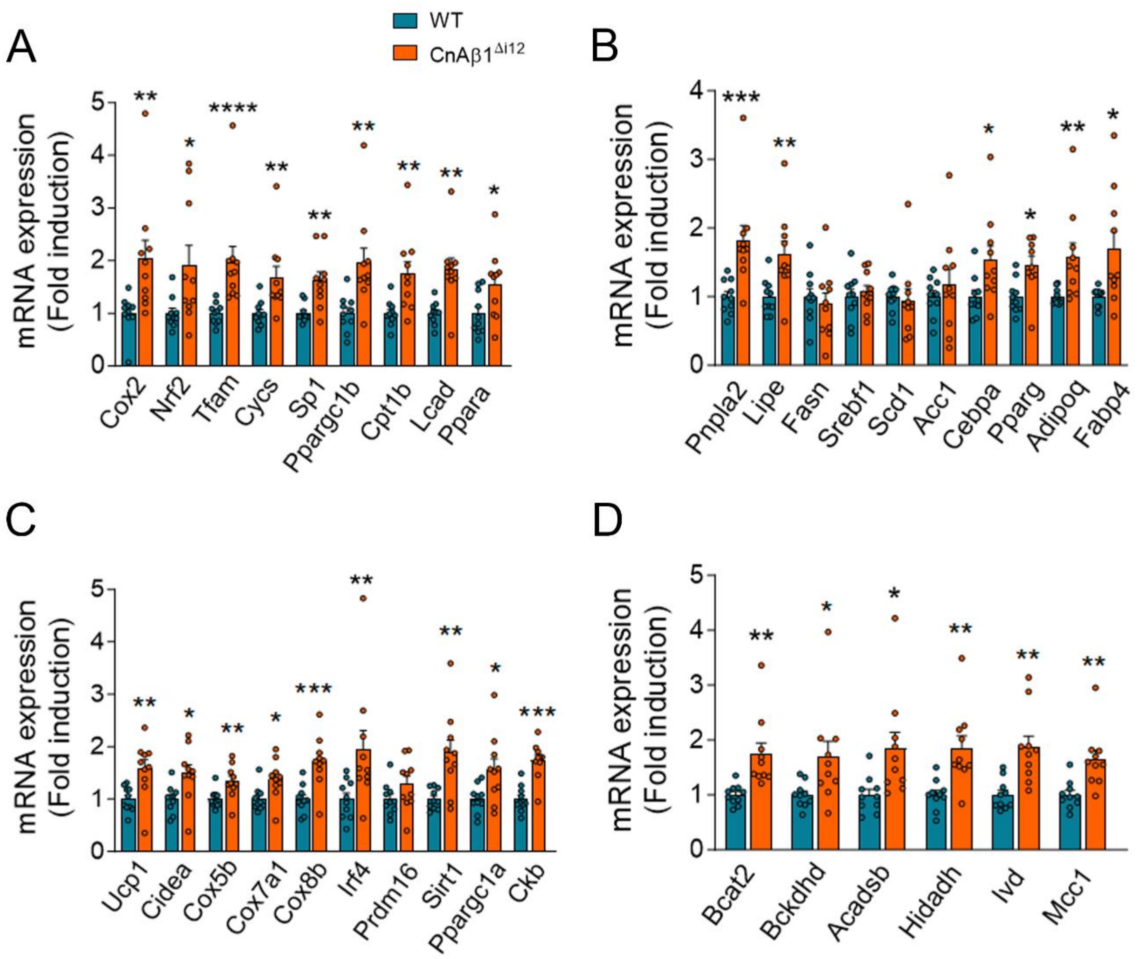


**Figure S3. CnAβ1$^{\Delta i12}$ mice on HFD show increased expression of mitochondria, lipolysis, fatty acid oxidation, thermogenic, and BCAA catabolism genes in BAT. A-D**, Gene expression related to mitochondria (Cox2, Nrf2, Tfam, Cycs, Sp1) and fatty acid oxidation (Ppargc1b, Cpt1b, Lcad, Ppara) (A), lipolysis (Pnpla2/Atgl, Lipe), lipogenesis (Fasn, Srebf1, Sced1, Acc1), and adipogenesis/differentiation (Cebp1, Pparg, Adipoq, Fabp4) (B), thermogenesis (C), and branched chain amino acid catabolism (D) was determined by qRT-PCR and normalized to β-actin mRNA in BAT of WT and CnAβ1$^{\Delta i12}$ mice after 6 weeks on HFD (n=10 per group). Results are expressed as mean ± SEM. *p<0.05; **p<0.01; ***p<0.001; ****p<0.0001 WT versus CnAβ1$^{\Delta i12}$ mice, unpaired Student's t-test and Mann-Whitney U test, depending on normality.

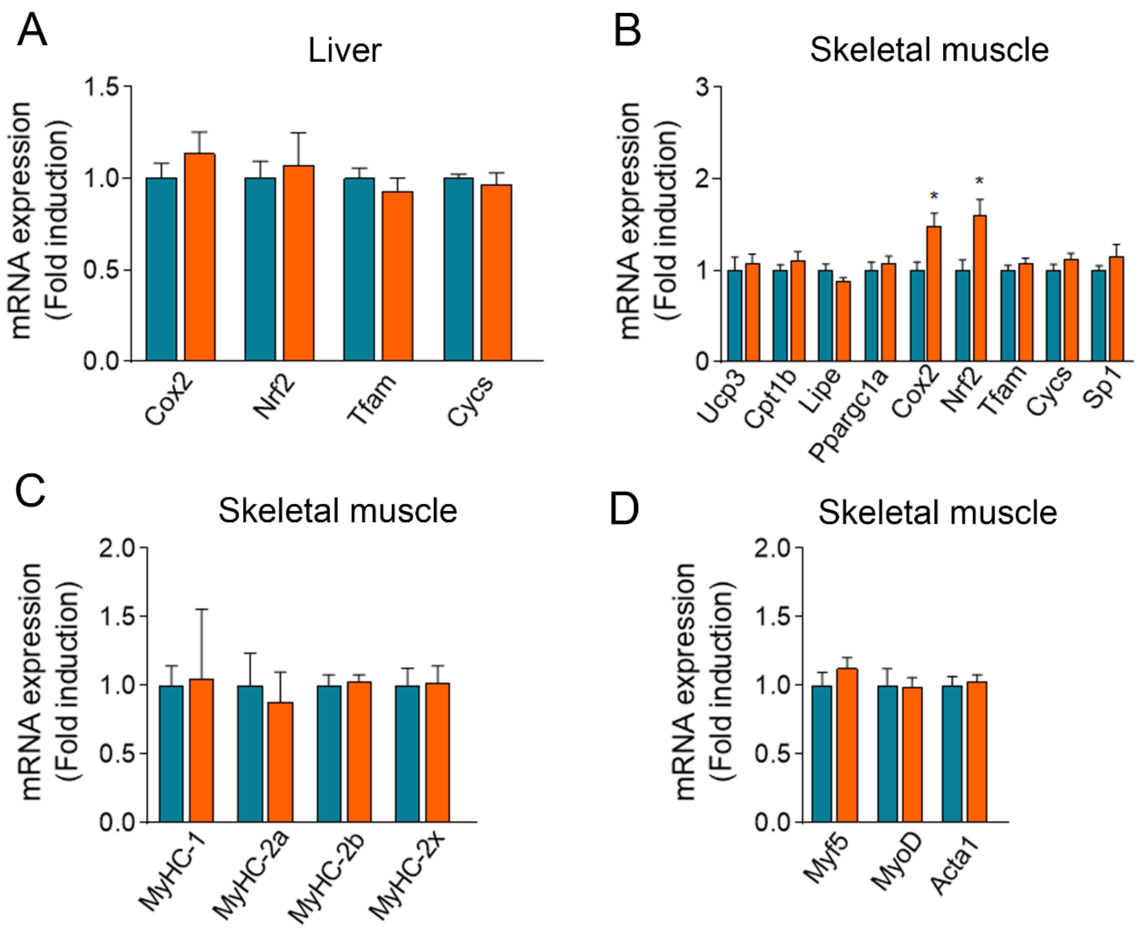


**Figure S4. Metabolic gene expression in liver and skeletal muscle of CnAβ1$^{\Delta i12}$ and WT mice. A**, Expression of mitochondria-related genes in the liver of WT and CnAβ1$^{\Delta i12}$ mice was measured by qRT-PCR after 6 weeks on HFD and normalized to β-actin. **B**, Expression of thermogenic and mitochondria-related genes in the tibialis anterior muscle was determined by qRT-PCR and normalized to β-actin mRNA in WT and CnAβ1$^{\Delta i12}$ mice after 6 weeks on HFD (n=10 per group). **C, D**, Expression of type I and type II skeletal muscle fibre genes (C) and skeletal muscle controlling transcription factors and fibres (D) was determined in the tibialis anterior by qRT-PCR and normalized to β-actin mRNA in WT and CnAβ1$^{\Delta i12}$ mice (n=9-10 per group) fed with HFD. No statistically significant differences were found in WT versus CnAβ1$^{\Delta i12}$ mice, unpaired Student's t-test and Mann-Whitney U test, depending on normality.
